# Non-canonical Telomerase Reverse Transcriptase Controls Osteogenic Reprogramming of Aortic Valve Cells Through STAT5

**DOI:** 10.1101/2022.08.23.504425

**Authors:** Rolando A. Cuevas, Luis Hortells, Claire C. Chu, Ryan Wong, Alex Crane, Camille Boufford, Cailyn Regan, William J. Moorhead, Michael J. Bashline, Aneesha Parwal, Angelina M. Parise, Parya Behzadi, Mark J. Brown, Aditi Gurkar, Dennis Bruemmer, John Sembrat, Ibrahim Sultan, Thomas G. Gleason, Marie Billaud, Cynthia St. Hilaire

## Abstract

**Background:** Calcific aortic valve disease (CAVD) is the pathological remodeling of valve leaflets. The initial steps in valve leaflet osteogenic reprogramming are not fully understood. As telomerase reverse transcriptase (TERT) overexpression primes mesenchymal stem cells to differentiate into osteoblasts, we investigated whether TERT contributes to the osteogenic reprogramming of valve interstitial cells (VICs).

**Methods:** Human control and CAVD aortic valve leaflets and patient-specific hVICs were used in in vivo and in vitro calcification assays. Loss of function experiments in hVICs and cells isolated from *Tert^-/-^*and *Terc^-/-^* mice were used for mechanistic studies. Calcification was assessed in *Tert^+/+^* and *Tert^-/-^* mice ex vivo and in vivo. In silico modeling, proximity ligation and co-immunoprecipitation assays defined novel TERT interacting partners. Chromatin immunoprecipitation and CUT&TAG sequencing defined protein-DNA interactions.

**Results:** TERT protein was highly expressed in calcified valve leaflets without changes in telomere length, DNA damage, or senescence markers, and these features were retained in isolated primary hVICs. *TERT* expression increased with osteogenic or inflammatory stimuli, and knock-down or genetic deletion of *TERT* prevented calcification in vitro and in vivo. Mechanistically, TERT was upregulated via NF-κB and required to initiate osteogenic reprogramming, independent of its canonical reverse transcriptase activity and the lncRNA *TERC*. TERT exerts non-canonical osteogenic functions via binding with Signal Transducer and Activator of Transcription 5 (STAT5). Depletion or inhibition of STAT5 prevented calcification. STAT5 was found to bind the promoter region of Runt-Related Transcription Factor 2 (*RUNX2*), the master regulator of osteogenic reprogramming. Lastly, we demonstrate that TERT and STAT5 are upregulated and colocalized in CAVD tissue compared to control tissue.

**Conclusions:** TERT’s non-canonical activity is required to initiate calcification. TERT is upregulated via inflammatory signaling pathways and partners with STAT5 to bind the *RUNX2* gene promoter. These data identify a novel mechanism and potential therapeutic target to decrease vascular calcification.

**Novelty and Significance:** *What is known?*

Calcific aortic valve disease (CAVD) is the most prevalent form of aortic valve pathology. CAVD strongly correlates with age and leads to heart failure and a high risk of stroke. Currently, the only therapeutic option is valve replacement, which comes with significant healthcare costs and additional risks to patients.

Runt-related transcription factor 2 (RUNX2) is the master transcription factor required for osteogenic differentiation of stem cells to osteoblasts and osteogenic reprogramming of cardiovascular cells. Yet, the early events driving its activity in aortic valve cells are poorly defined.

In addition to its reverse transcriptase enzymatic activity, TERT exhibits non-canonical transcriptional regulatory functions and overexpression of TERT primes mesenchymal stem cells to differentiate down the osteoblast lineage.

*What new information does this article contribute?*

TERT protein levels in calcified aortic leaflets and valve interstitial cells, and its non-canonical osteogenic activity are independent of changes in telomere length and cell senescence.

Genetic loss or depletion of TERT prevented calcification in valve interstitial cells, coronary smooth muscle cells, and mesenchymal stem cells in vitro and the vasculature in vivo.

Early in the osteogenic reprogramming inflammatory signaling promotes TERT to co-localize with SMARCA4 and STAT5, and this TERT-tethered STAT5 binds to the *RUNX2* gene promoter, the master regulator of osteogenic transcriptional programs.

STAT5 depletion and pharmacological inhibition prevent calcification of human valve interstitial cells, coronary smooth muscle cells, and mesenchymal stem cells.

*What are the clinical implications?*

We have identified TERT-STAT5 as a novel signaling axis that orchestrates the early steps in the osteogenic reprogramming of aortic valve cells. Inhibiting TERT/STAT5 interaction or their activity may be leveraged for the development of therapeutic strategies to halt or prevent calcification in the aortic valve, bioprosthetic valves, and or perhaps other cardiovascular tissues.

Invasive and expensive surgical procedures are currently the only treatment option for patients with CAVD. The discovery and defining of the early events driving vascular calcification identifies novel and druggable targets for developing non-surgical therapies.

## INTRODUCTION

Calcific aortic valve disease (CAVD) encompasses a spectrum of pathological remodeling of the valve leaflet, ranging from mild valve thickening that displays microcalcifications to macrocalcifications that predominate in more advanced disease stages.^1,2^ The advanced state, aortic stenosis, results from fibrotic remodeling and the formation of calcified nodules on the valve leaflet that impede and disrupt blood flow, causing excessive strain on the cardiac tissue, increasing the risk of stroke and leading to heart failure.^3^ CAVD severity and incidence increase with age, with CAVD prevalence >1,000 per 100,000 individuals ≥75 years of age.^4,5^ Further, roughly 0.4-0.6% of the population harbors bicuspid aortic valve malformations, and these malformed valves are prone to calcify and develop CAVD.^4^ Currently, no therapeutic treatment halts or reverses pathologic calcification that occurs in the aortic valve leaflets or other cardiovascular tissues. The only therapeutic option to treat aortic stenosis is valve replacement via surgical or transcatheter procedures, incurring substantial medical costs and health care burdens to the patient.^6^

Aortic valve interstitial cells (VICs) are the primary cell type within the valve leaflet and are responsible for maintaining leaflet integrity. VICs reside in a quiescent state; however, adaptive or maladaptive responses to environmental factors such as inflammation, extracellular matrix (ECM) breakdown, and mechanical stress may disrupt VIC homeostasis.^7,8^ With alterations in homeostasis, VICs acquire a myofibroblast-like phenotype, termed “activated VICs,” capable of proliferation, contraction, and secretion of proteins that further remodel the extracellular milieu.^9^ Activated VICs can transition into calcifying cells; however, the initial steps regulating this process are not fully understood. It is well established that the osteogenic reprogramming of cardiovascular cells such as VICs and vascular smooth muscle cells (SMCs) is analogous to the differentiation of mesenchymal stem cells (MSCs) into bone-forming osteoblasts; osteogenic differentiation of MSCs and the reprogramming of cardiovascular cells is primarily orchestrated by Runt-related transcription factor 2 (*RUNX2*), the master regulator of osteogenic transcriptional programs.^10–12^ Other osteogenic signature markers, such as osteocalcin, alkaline phosphatase, and osteopontin, are upregulated in the differentiation of stem cells to osteoblasts and the calcifying of VICs.^9,13^ While it is accepted that VICs upregulate the expression of osteogenic genes in response to stresses such as inflammation, ECM remodeling, and mechanical stress, the specific mechanistic steps between those stresses and the initial reprogramming events in the activation of osteogenic transcriptional reprogramming remain ill-defined.

The telomerase complex extends and maintains the ends of chromosomes after cell division. The core complex consists of the telomerase long non-coding RNA component (*lncTERC*) and telomerase reverse transcriptase (TERT) protein, along with other scaffolding proteins.^14^ In addition to reverse transcriptase activity, TERT also exhibits non-canonical functions independent of telomere extension, such as regulating gene expression and orchestrating chromatin remodeling.^15–17^ For example, TERT acts as a cofactor in Wnt-dependent promoter activation by interacting with the chromatin remodeling protein SWI/SNF related, matrix associated, actin dependent regulator of chromatin, subfamily a, member 4 (SMARCA4), facilitating the expression of Wnt/β-catenin target genes in stem cells.^18^ In a murine model of atherosclerosis, TERT induced chromatin remodeling and SMC proliferation by chaperoning the retinoblastoma-associated transcription factor 1 (E2F1) binding to S-phase gene promoters.^19,20^ Linking together the themes of calcification and non-canonical TERT, two co-published studies identified that the overexpression of *TERT* primed human MSCs to differentiate down the osteogenic lineage and develop bone-like structures.^21,22^ The mechanism by which TERT drives the osteogenic differentiation of human MSCs is unknown. Collectively, these studies provide strong evidence of telomerase-independent functions of TERT in altering chromatin states and regulating gene expression via various protein-protein interactions.

Although telomere length is recognized as a biomarker of aging, it can vary significantly among individuals of the same age, including twins.^23^ Studies have examined the association between shortened telomeres in circulating leukocytes and various vascular conditions, with mixed results.^24–26^ One study compared telomere lengths in leukocytes of control patients and those with atherosclerotic plaques, as well as the atherosclerotic plaques themselves and found that while atherosclerotic patients’ leukocytes had shorter telomeres than controls, their plaque cells exhibited longer telomeres than both groups’ leukocytes, suggesting heightened telomerase activity in the diseased tissue.^27–29^ Thus, whether leukocyte telomere length can be used as a biomarker remains speculative. A notable aspect of these prior studies is the lack of TERT protein assessment or a focused investigation of telomere length in disease tissues.

Signal transducers and activators of transcription (STAT) proteins play essential roles in inflammatory, proliferative, and apoptotic programs, mediating the signal transduction pathways of various cytokines and growth factors.^30^ STAT proteins function as transcription factors and recruit chromatin remodeling and histone-modifying proteins to upregulate gene transcription.^31^ STAT5A and STAT5B are encoded by two highly conserved genes in humans and mice; 96% of their amino acid sequence is shared, and many of their activities are redundant.^32^ Herein, unless specific tools are utilized to allow a distinction, we will generally refer to both as STAT5. Upon phosphorylation, STAT5 dimerizes and translocates into the nucleus to bind the consensus motif (TTCNNNGAA) and activates the promoters of various target genes by recruiting transcriptional co-activators.^33,34^ Crucially, studies have shown that several osteogenic genes depend on STAT5 to promote chondrocyte and osteoblast differentiation.^35,36^

In this study, we tested the hypothesis that TERT contributes to the osteogenic reprogramming of VICs. We utilized primary human aortic valve tissues from control donors and CAVD patients and generated patient-specific VIC lines for in vitro disease modeling. We established baseline patterns and osteogenic induction of *TERT* and calcification markers in CAVD tissue and VICs. We assessed the consequences of genetic deletion or depletion of *TERT* during osteogenic reprogramming in several cell types and in in vivo and ex vivo calcification models. We further defined the underlying mechanism by which TERT and STAT5 participate in the osteogenic reprogramming of VICs.

## METHODS

### Data Availability

The Materials, Methods, and Major Resources Table are found in the Supplemental Material. The raw data, experimental materials, and analytic methods are available from the corresponding author upon reasonable request.

## RESULTS

### TERT is elevated in CAVD tissues

Human aortic valves were collected after surgical aortic valve replacement procedures or from cadaveric tissue obtained via the Center for Organ Recovery and Education (CORE) of the University of Pittsburgh Medical Center and processed as previously described.^37^ Macroscopic examination and histological staining with Von Kossa were used to determine and confirm whether valves could be classified as a non-calcified control or as having calcific aortic valve disease (CAVD) (**Figure 1A, Supplemental Figure 1A, Supplemental Table 1**). RUNX2 is the master transcription factor required to initiate stem cells to differentiate into osteoblasts and is critical for the pathological osteogenic reprogramming of vascular cells.^38–41^ Immunofluorescent staining showed RUNX2, and the late calcification marker osteopontin (OPN) were significantly upregulated in CAVD tissue compared to control tissues, indicating activation of osteogenic transcriptional programs in CAVD tissues (**Supplemental Figure 1B**). Previous studies found that overexpression of TERT primes human MSCs to differentiate down the osteogenic lineage.^21,22^ We found that *TERT* transcript levels were significantly upregulated in CAVD tissue relative to control samples (**Figure 1B**), and *TERT* expression levels showed no correlation with the donor’s age (**Supplemental Figure 1C**). Markers of senescence (cyclin-dependent kinase inhibitor 1A, *CDKN1A*, galactosidase beta 1, *GLB1*), proliferation (proliferating cell nuclear antigen, *PCNA*), DNA damage (tumor protein 53, *TP53*), and cytoskeletal markers (actin alpha 2 smooth muscle, *ACTA2*, vimentin, *VIM,* interleukin 6, *IL6*) showed no statistically significant differences, while the inflammatory marker and tumor necrosis factor-alpha (*TNF)* was significantly elevated in CAVD samples (**Supplemental Figure 1D**), supporting observations by others that indicate a role for inflammatory signaling in aortic valve calcification pathogenesis.^42,43^

**Figure 1.**
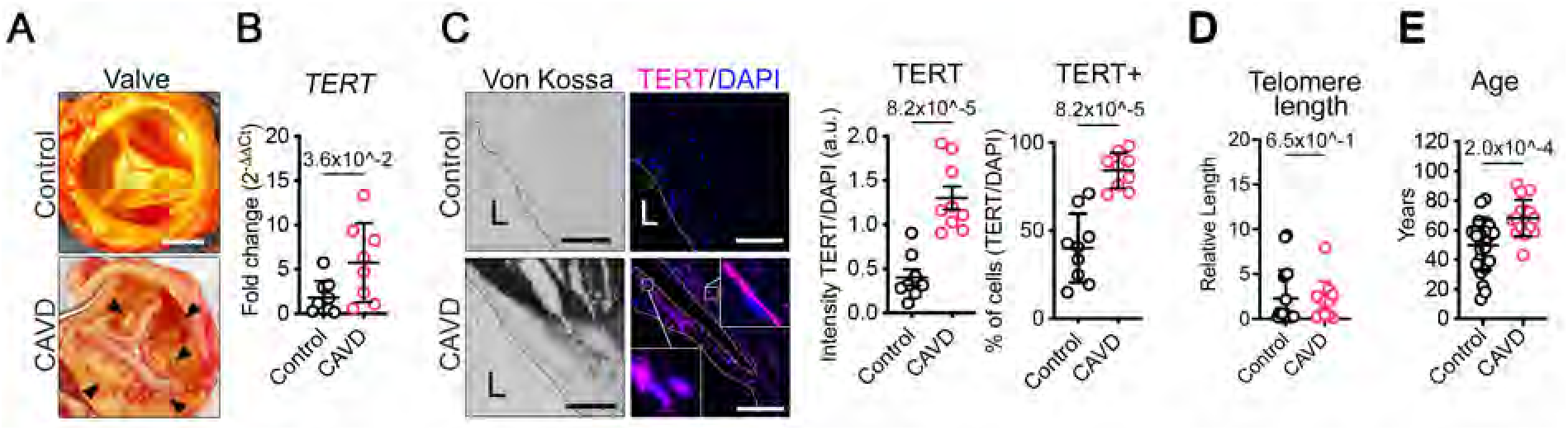
TERT is elevated in CAVD tissues. (A) Examples of non-calcified control (top) and calcified CAVD (bottom) human valves used in this study. Arrowheads indicate calcified nodules Scalebar. 1 cm. (B), TERT mRNA transcript quantification in control and CAVD valve tissues. N = 8 control, n= 9 CAVD. (C), Representative serial sections of Von Kossa (dark precipitation) and TERT immunofluorescent staining in control and CAVD valve tissues. Scalebar 100 μm. Graphs show the quantification of TERT intensity and the number of TERT-positive cells. n = 9 control, n = 9 CAVD. (D), Telomere length measurements in valve tissues. n = 19 control, n = 12 CAVD. (E), Age of the patients utilized in this study. n = 25 control, n = 16 CAVD. Data are shown as means ± SD. P values were calculated using the Mann-Whitney U test and shown on each graph. L = lumen.

We detected elevated TERT protein signal in CAVD valves compared to non-calcified controls, with the signal localized to areas of calcification. Similarly, the number of cells expressing TERT was significantly elevated in our CAVD tissues compared to non-calcified control tissues (**Figure 1C**). No statistically significant differences were observed in the staining levels of proliferation marker PCNA or the DNA damage indicator phosphorylated gamma histone 2AX (γ-H2AX) between CAVD and control tissues (**Supplemental Figure 1E**). We also found no statistically significant differences in the relative mean telomere length between the control and CAVD tissues in our study (**Figure 1D**), indicating no overt influence of the canonical telomere-extending functions of TERT. The mean age of CAVD patients in this study was slightly elevated compared to the control patients, 68 and 54 years old, respectively (**Figure 1E**). Together, these data show that TERT, osteogenic markers, and inflammatory signatures are upregulated in CAVD tissues compared to controls, while markers indicating alterations in proliferation, DNA damage, and senescence are not altered, suggesting that the non-canonical activity of TERT is operative.

### CAVD human VICs at baseline recapitulate observations in CAVD tissues

We created patient-specific valve interstitial cell (VIC) lines from CAVD and control valves.^37^ At baseline, freshly isolated hVICs from CAVD and control patients exhibited no morphological or statistical differences in their proliferative or migratory capacity (**Figure 2, A-C, and Supplemental Figure 1F**). TERT and RUNX2 proteins were significantly elevated at baseline in CAVD hVICs compared to control hVICs (**Figure 2D**). We observed no statistical differences in the protein levels of the hVIC activation markers SM22 or αSMA between CAVD and control hVICs, but observed a significant increase of the mineralizing enzyme tissue non-specific alkaline phosphatase (TNAP) in CAVD hVICs at baseline (**Supplemental 1G**). As in the leaflets, there was no statistical difference in the relative telomere length between CAVD and control hVICs (**Figure 2E**). These data show that hVICs isolated from CAVD patients display gene expression and protein signatures that recapitulate the osteogenic phenotype observed in CAVD leaflet tissue.

**Figure 2.**
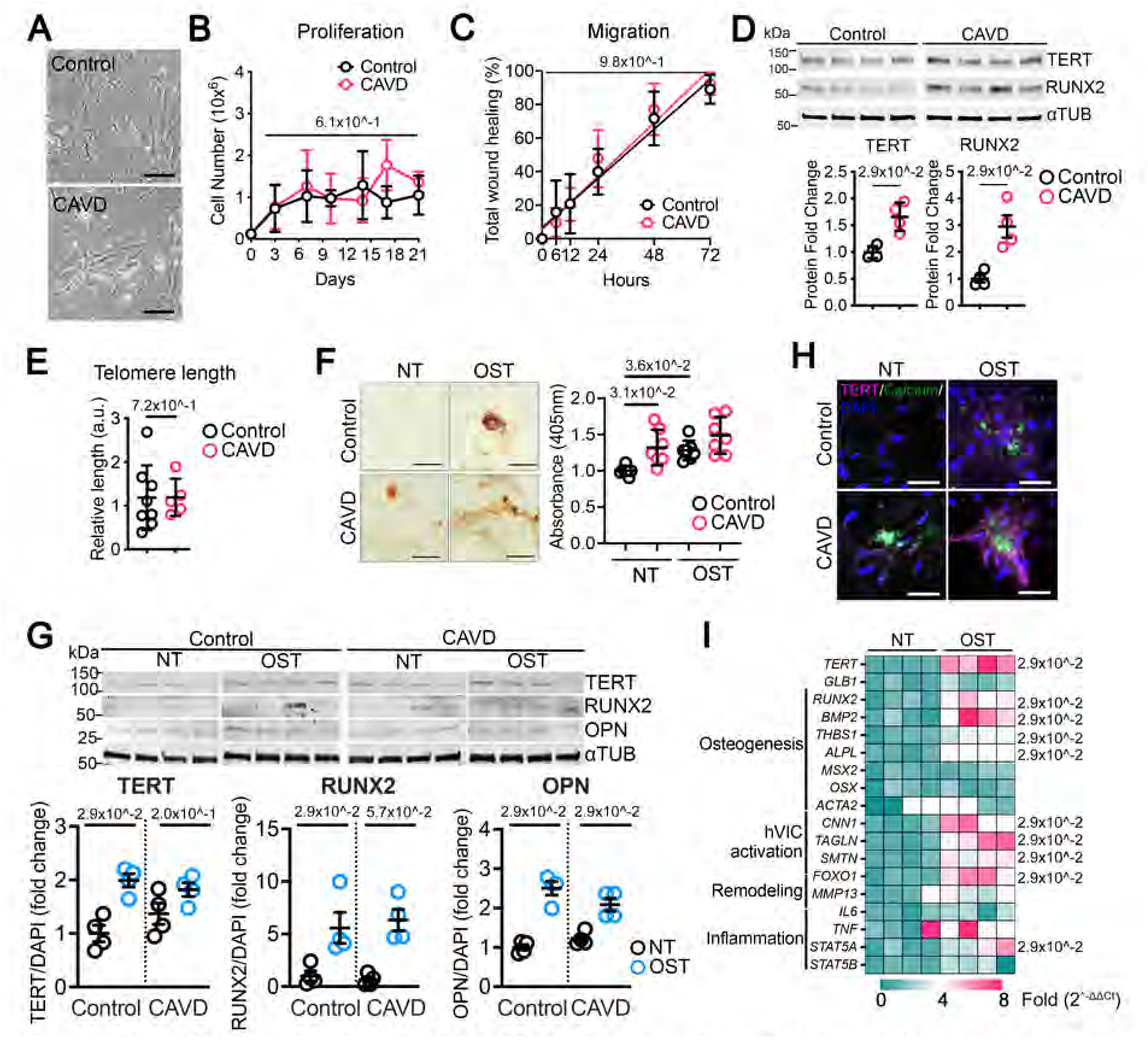
TERT is elevated in CAVD hVICs at baseline and upregulation during osteogenic reprogramming. (A), Representative images showing cell morphology of control and CAVD hVICs. Scalebar 50 μm. (B), Proliferation of control and CAVD hVICs at baseline. n = 3 both groups. (C) Relative migration distances of control and CAVD VICs. n = 10 both groups. (D) Western blot staining of control and CAVD hVICs at baseline. Quantification of protein levels is shown on the right panels. n = 4 both groups. (E) Telomere length measurements in VICs. n = 8, control, n = 5 CAVD. (F) Representative images of early calcification in hVICs growing in normal conditions (no treatment NT) or stimulated with osteogenic media (OST) for 14 days. Calcium deposition was visualized by Alizarin Red staining. Scalebar 50 μm. Quantification of calcification on the right graphs. n = 7 both groups. (G), Western blot staining of samples from control and CAVD hVICs after 14 days of NT or OST. Quantification of protein levels is shown on the bottom graphs. n = 4 both groups. (H), Representative immunofluorescent staining images of TERT in control and CAVD hVICs after 14 days of NT or OST. Calcium deposition was visualized with OsteoImage. Scalebar 100 μm. (I) Expression profile control VICs after 14 days of NT and OST treatment. n = 4 both groups. Data are shown as means ± SD. P values were calculated with the Mann-Whitney U test (Figures B, C, D, E, G, and I) and the Kruskal-Wallis H test with Dunn post hoc test (Figure F) and shown on each graph

### TERT expression is upregulated in human VICs in an in vitro osteogenic disease model

To model CAVD in vitro and investigate a role for TERT during hVIC osteogenic reprogramming, we cultured CAVD and control hVICs in osteogenic media conditions (OST), a media widely utilized to induce osteogenic differentiation of stem cells to osteoblasts.^44^ Alizarin Red staining for calcification revealed that both OST-treated CAVD and control hVICs lay down calcified matrix as early as 14 days (d14), CAVD hVICs calcified earlier relative to control hVICs (**Figure 2F**). Importantly, we observed that CAVD hVICs calcify de novo under no treatment (NT) conditions, suggesting that CAVD hVICs are primed for osteogenic reprogramming (**Figure 2F**). The calcification markers RUNX2 and OPN were significantly upregulated in both control and CAVD hVICs at d14 OST, indicating active osteogenic reprogramming (**Figure 2G**). TERT protein levels were also significantly increased in control and CAVD hVICs after d14 of OST (**Figure 2G**). TERT protein signal can be seen in CAVD hVICs at d14 under NT conditions (**Figure 2H**, left panels). At d14 of OST, the TERT protein staining was elevated and localized near areas of calcification in both control and CAVD hVICs; however, CAVD hVICs formed larger calcified nodules (**Figure 2H**, right panels). We observed no significant differences in cell numbers at the end of the d14 assay under NT or OST treatment conditions (**Supplemental Figure 2A**). As we observed that CAVD hVICs exhibit elevated osteogenic markers at baseline (**Supplemental Figure 1G**) and are prone to calcify (**Figure 2H**) de novo, we assessed transcriptional differences between control hVICs under NT or OST. Osteogenic markers *BMP2*, *THBS1*, *ALPL,* and myofibroblast markers *CNN1*, *TAGLN*, and *SMTN* were significantly upregulated in control hVICs at d14 of OST treatment (**Figure 2I**). Likewise, the calcification-associated gene *FOXO1* and the CAVD-associated metalloproteinase *MMP13* were also significantly upregulated in OST conditions.^45,46^ Lastly, the pro-inflammatory transcription factor signal transducer and activator of transcription 5A *(STAT5A)* was significantly upregulated, while the expression of the isoform *STAT5B* remained unchanged. These results show that hVICs are undergoing osteogenic reprogramming and not de novo mineral nucleation.^47^

In cancer and stem cells, elevated expression of TERT is associated with long telomeres and enables unlimited proliferation, while short telomeres can trigger cellular senescence.^48,49^ Although we observed no differences in telomere length in our tissues and cells in this study (Figures 1D and 2E), we assessed cellular senescence and observed no significant differences in the expression of senescence-associated β-galactosidase (SA-β-gal, *GLB1*) between CAVD and control hVICs at baseline **(Supplemental Figure 2B)** nor increased in control hVICs in OST treatment (**Figure 2I**). SA-β-gal activity staining indicated that cellular senescence was not engaged even after 28 days of OST treatment (**Supplemental Figure 2C**). These data confirmed that cellular senescence is not operative during hVICs osteogenic reprogramming.

NF-κB is a transcription factor complex that acts downstream of inflammatory signals to mediate inflammatory responses, and the c-Myc transcription factor mediates cell proliferation and differentiation. Others have shown that both NF-κB and c-Myc can upregulate transcription of the *TERT* gene.^50–52^ We found that inhibiting NF-κB with BAY-107082, an irreversible inhibitor that blocks the phosphorylation of IκB-α,^53,54^ inhibited *TERT* expression during the osteogenic reprogramming of hVICs. We also observed downregulation of *RUNX2* expression, consistent with upstream TERT’s regulatory role on *RUNX2* expression (**Supplemental Figure 2D**). Likewise, we found that inhibiting c-MYC with 10058-F4 inhibited *TERT* expression during the osteogenic differentiation of hVICs (**Supplemental Figure 2E**), validating previous research showing that c-MYC regulates *TERT* gene expression.^16^ These results further support that inflammatory signaling contributes to the development of CAVD.

### TERT is required for the osteogenic differentiation of human MSCs

Previous studies have shown that the overexpression of TERT primes human mesenchymal stem cells (hMSCs) to differentiate into osteoblasts.^21,22^ To assess the role of endogenous TERT in osteogenic differentiation, we subjected hMSCs to NT and OST treatment. OST treatment induced robust calcification as early as d14, while cells in NT conditions did not calcify (**Supplemental Figure 3A**). Protein analysis showed that the TERT protein levels significantly increased on day 3 of the OST treatment, coinciding with increased *RUNX2* expression and RUNX2 protein levels (**Supplemental Figures 3B** and **3C**). hMSCs in OST treatment exhibited intense TERT staining, and TERT-positive cells clustered around calcified nodules (**Supplemental Figure 3D**). Using lentiviral-mediated transduction of expressing short-hairpin RNA to deplete TERT (shTERT) or scrambled control (shControl), we found that knockdown of *TERT* significantly inhibited the calcification of hMSCs (**Supplemental Figures 3E** and **3F**). While previous studies show that overexpression of *TERT* primed hMSCs to undergo osteogenic differentiation, our data show that TERT is required for the osteogenic differentiation of hMSCs.^21,22^

**Figure 3.**
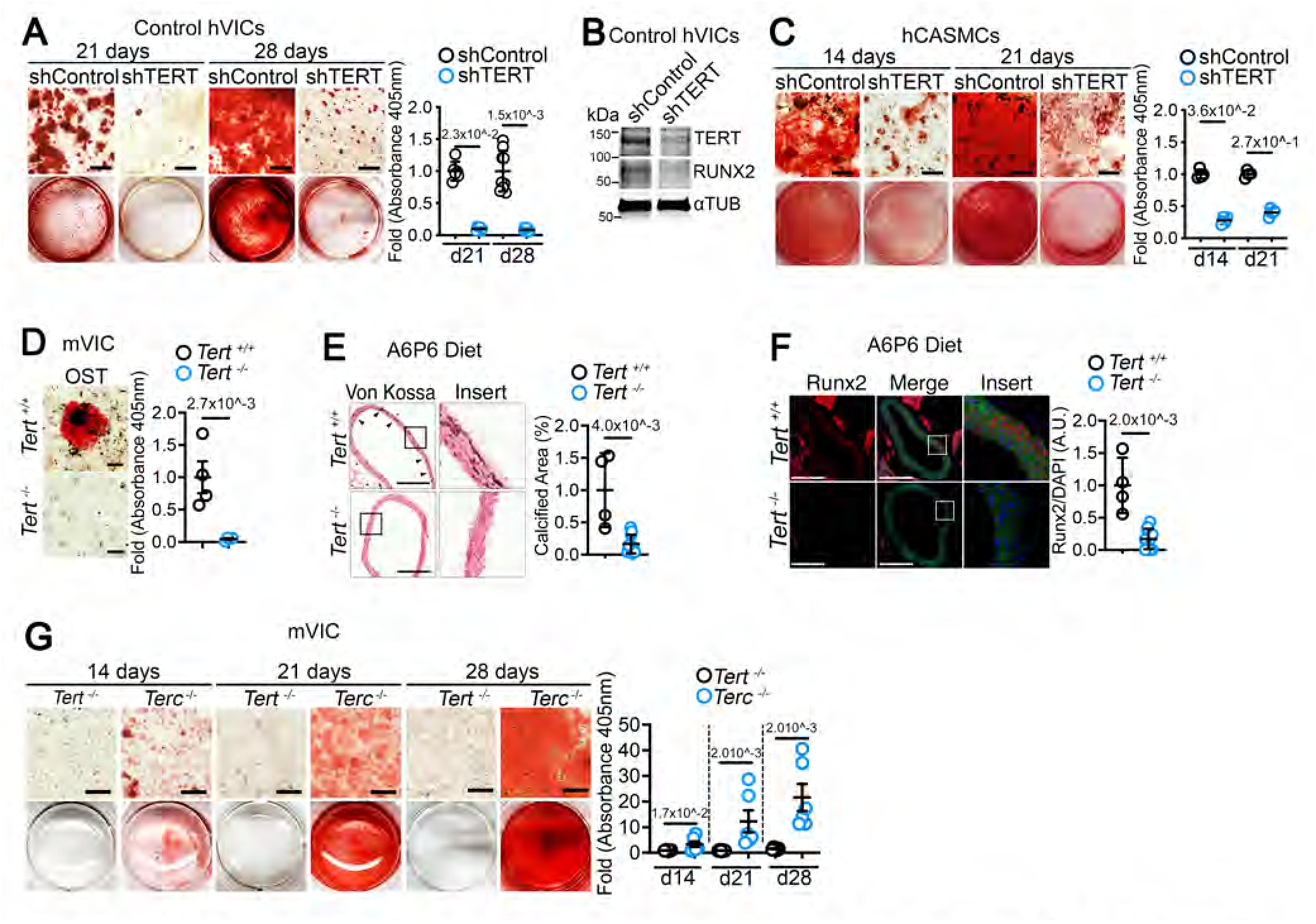
TERT is required to initiate osteogenic transition and calcification. (A) Control hVICs were transduced lentivirus containing shControl or shTERT and followed by OST stimulation for up to 28 days. Calcium deposition was visualized by Alizarin Red staining. n = 6 shControl, n = 6 shTERT at 21 days and n = 9 each group at day 28. Scalebar 400 μm. (B) Representative western blot staining of control hVICs transduced with lentivirus containing either shControl or shTERT followed by 7 days of OST. (C) hCASMCs were transduced with shControl or shTERT and stimulated with OST for 14 and 21 days. n = 4 for each group at both time points. Scalebar 400 μm. (D) Representative images of mVIC isolated from *Tert^+/+^* and *Tert^-/-^* mice were stimulated with OST for 28 days. Scalebar 400 μm. (E) Representative Von Kossa staining images of *Tert^+/+^* and *Tert^-/-^* mice aortas (dark precipitation). Scale bar 400 μm. Quantification of calcification is shown on the right panel. (F) Representative immunofluorescent images of Runx2 expression indicative of calcification (red, insert) on *Tert^+/+^* and *Tert^-/-^* mice aorta. Scale bar 400 µm. Runx2 quantification of calcification is shown on the right panel. (G) Representative images of mVICs isolated from F1 *Tert^-/-^ or Terc^-/-^* mice were stimulated with OST for up to 21 days. Scalebar 100 μm. Data are shown as means ± SD. P values were calculated using the Kruskal-Wallis H test, Dunn post hoc test (Figures A, C, and G), and Mann-Whitney U test (Figures D, E, and F).

### TERT is required for calcification in vitro and in vivo

Control hVICs were transduced with shControl and shTERT lentivirus, and like hMSCs, knockdown of *TERT* significantly inhibited calcification compared to the control group and downregulated RUNX2 protein levels (**Figures 3A** and **3B**). We assessed TERT’s role in calcifying human coronary artery smooth muscle cells (hCASMCs), which calcify in atherosclerotic plaques.^55^ Knocking down TERT significantly inhibited hCASMCs calcification (**Figure 3C**). Together, these data indicate that TERT is essential for the osteogenic reprogramming of hVICs and hCASMCs. Next, mice VIC (mVICs) and bone marrow MSCs (mBMMSCs) lines were generated from *Tert^+/+^* and *Tert^-/-^* mice.^56^ In line with our observations in human cells with depleted TERT, we observed significant inhibition of calcification in *Tert^-/-^* mVICs (**Figure 3D**) and *Tert^-/-^* mBMMSCs (**Supplemental Figure 4A**) compared to *Tert^+/+^* cells. No statistically significant differences were observed in senescence-associated β-galactosidase activity in mVICs or mBMMSCs (**Supplemental Figures 4B** and **4C**).

**Figure 4.**
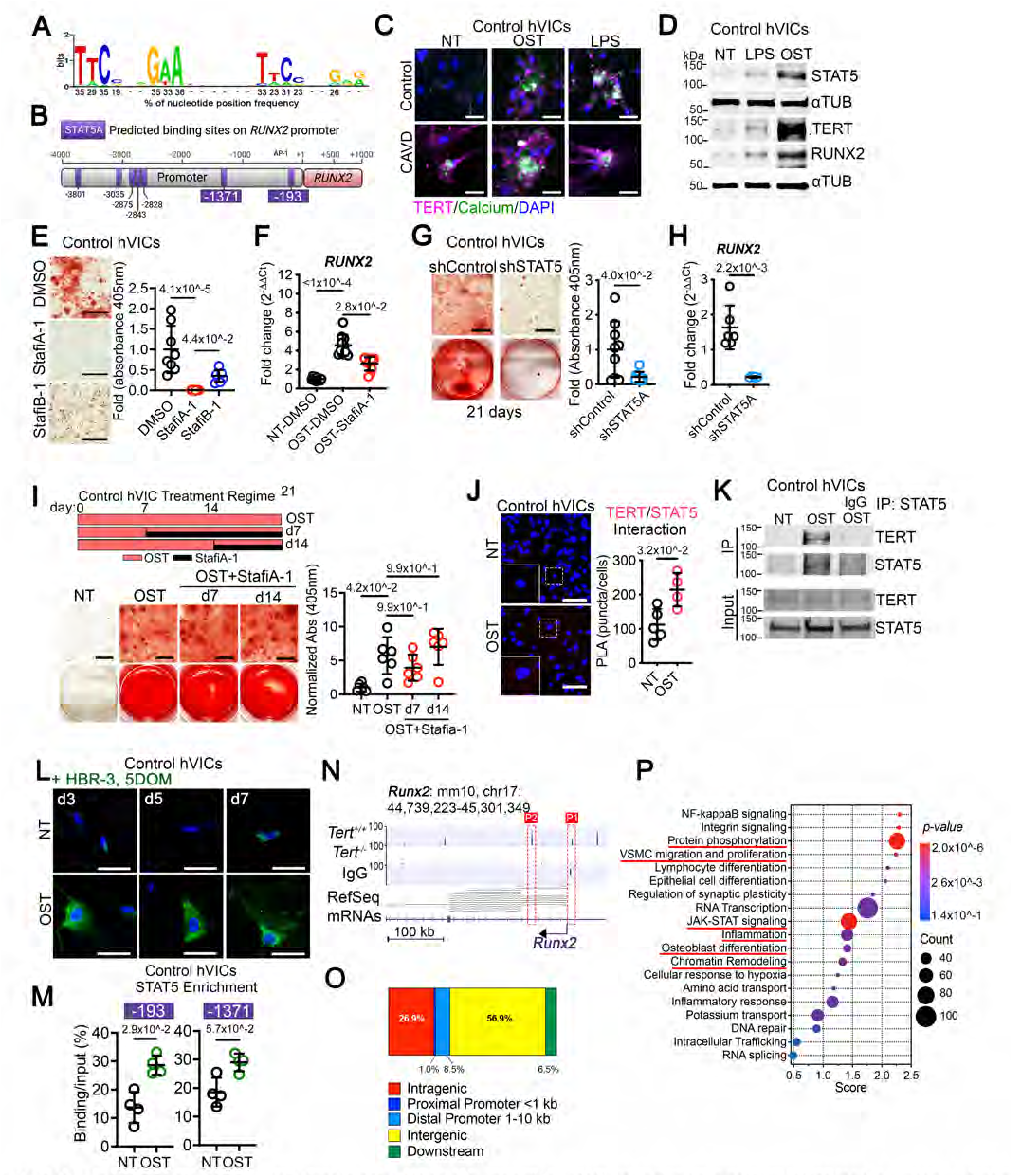
STAT5 binds to TERT and is required for the calcification of human VICs. (A) Graphic of in silico analysis identifying the consensus binding site for STAT5. (B), Diagram depicting STAT5 binding to RUNX2 promoter and the positions of the predicted STAT5 binding sites (purple boxes). (C) Representative immunofluorescence images of control hVICs after 14 days of LPS treatment or OST stimulation. Scalebar 50 μm. (D) Representative western blot staining of hVICs after 14 days of OST stimulation. (E) Representative images of control hVICs treated with 10 μM of the STAT5 inhibitors StafiA-1 or StafiB-1 throughout 28 days of osteogenic stimulation. Scalebar 400 μm. n = 8 each group. The quantification of calcification is shown on the right graph. (F) RUNX2 mRNA transcript quantification at d7 OST, n = 10. (G) Control hVICs were transduced with lentivirus containing shControl or shSTAT5 constructs and treated with OST for 21 days. Calcium deposition was visualized by Alizarin Red staining. n = 9 for each group. Scalebar 400 μm. (H) RUNX2 mRNA transcript quantification at d7 OST, n = 6. (I) Schematic of OST treatment with delayed shTERT transduction (top panel).Representative images of hVIC (left panels) treated with 10 μM of the STAT5 inhibitors StafiA-1 at d7 or d14 of OST treatment. Samples were collected at d21 (bottom panel). Scalebar 400 μm. n = 6 for each group. (J) Representative images of TERT/STAT5 complex (red foci) in control hVICs cultured for 21 days in osteogenic medium and detected by PLA. Scalebar 100 μm. n = 4 for each group. (K) STAT5 co-immunoprecipitation in control hVICs under OST for 10 days to detect TERT/STAT5 interaction. (L) Control hVIC co-expressing TERT-NFAST and STAT5A-CFAST were labeled with 5 μM HBR-3,5DOM (4-hydroxy-3,5-dimethoxybenzylidene rhodamine, coral) and imaged at d3, d5, and d7 of osteogenic treatment (OST) or control no-treatment (NT). Scale bars 50 μm. (M) Chromatin immunoprecipitation shows enrichment of STAT5 on RUNX2 promoter OST conditions compared to NT. The base position is indicated in the blue boxes. n=4 for each group as in B. (N) STAT5 CUT&Tag sequencing tracks in genomic regions of Runx2 in no-treatment control and osteogenic condition-treated mice VICs. (O) Stacked chart representing genome-wide STAT5 occupancy in promoters, genes, and intergenic regions frequency in mice VICs. (P) Top 20 GO pathways enriched in genes with significant loss of STAT5 in Tert+/+ vs Tert-/-mice VICs. Data are shown as means ± SD. P values were calculated by the Mann-Whitney U test (Figures F, H, and J) and the Kruskal-Wallis H test (Figures E and G) and indicated on top of each graph.

To determine whether TERT is required to initiate and/or maintain osteogenic reprogramming, we cultured control hVICs in OST and transduced shTERT lentivirus on d7 or d14, and calcification was assessed on d21. Control hVICs transduced with shTERT on day 7 showed a significant reduction in calcification compared to the OST alone, while transduction of shTERT on day 14 did not show any decrease in calcification (**Supplemental Figure 4D**), indicating that TERT is required to initiate, not maintain, reprogramming to the osteogenic phenotype.

Next, we used *Tert^+/+^* and *Tert^-/-^* mice strains to test whether TERT is required for diet-induced non-atherosclerotic medial arterial calcification in vivo in mice. Of note, *Tert^-/-^* mice were generated from heterozygous breeding pairs (F1 generation) and thus did not exhibit shortened telomeres.^57^ *Tert^+/+^* and *Tert^-/-^* mice were fed an adenine-phosphate (A6P6) diet, which was previously shown to induce medial arterial calcification in the vasculature of mice.^58^ At the end of the 12-week regime, we observed significantly higher levels of calcification in the aorta of *Tert^+/+^* compared to *Tert^-/-^* mice (**Figure 3E**). Staining for Runx2 protein mirrored calcification presentation and was significantly higher in the *Tert^+/+^* compared to *Tert^-/-^* tissue (**Figure 3F**). These findings were validated in an ex vivo calcification assay. Thoracic aortic rings obtained from the *Tert^+/+^* and *Tert^-/-^* mice were cultured in ex vivo OST or control NT medium. *Tert^+/+^* aortic rings exhibited significantly elevated calcification compared to aortic rings from *Tert^-/-^* mice on d14 (**Supplemental Figure 4E**). *Tert^+/+^* aortic rings also exhibited prominent staining for Runx2 protein, which was absent in aortic rings from *Tert^-/-^* mice (**Supplemental Figure 4F**), and western blot analysis confirmed elevated Runx2 protein levels in *Tert^+/+^* aortic rings compared to *Tert^-/-^* (**Supplemental Figure 4G**). Neither calcification nor Runx2 was detected in the NT groups. Together, these data show that TERT is required for osteogenic reprogramming of cells in vitro and in vivo.

TERT’s canonical telomere-extending activity requires the telomerase long non-coding RNA component (*lncTerc*) to maintain chromosome telomere length and genome integrity.^14,59^ *LncTerc* exerts a dual function in the telomerase complex; it guides the telomerase complex to the telomere ends and is the primer for TERT reverse transcriptase activity, and it serves as a scaffold for tethering additional proteins to the telomerase complex (i.e., Dyskerin, NOP10, GAR1, and NHP2).^60^ To evaluate that TERT’s role in calcification is independent of its canonical telomere-extending function, we isolated mVICs from *Terc*^-/-^ mice. While *Tert^-/-^* mVICs did not calcify under OST conditions, *Terc^-/-^* mVICs were significantly calcified (**Figure 3G**). Similarly, in hVICs, BIBR1532, which blocks the reverse transcriptase activity of TERT,^61^ did not prevent calcification nor decrease the expression of *RUNX2* (**Supplemental Figure 4H** and **4I**). Together, these data illustrate that the canonical telomere-extending function of TERT does not contribute to osteogenic reprogramming.

### TERT/STAT5 interact and bind to the *RUNX2* gene promoter to initiate osteogenic reprogramming

Several studies have shown that TERT participates in gene transcription by physically interacting with transcription factors.^17,20,62,63^ RUNX2 is the master regulator of osteogenic differentiation of stem cells.^38,39^ Increased *RUNX2* expression and transcriptional activity are the hallmarks of osteoblast differentiation and cardiovascular calcification and are required for osteogenic reprogramming of vascular cells.^40,41^ As our data show that RUNX2 is significantly downregulated in *TERT*-knockdown and deficient cells (Figure 3), we used the LASAGNA^64^ and TRANSFAC/MATCH^65^ programs to identify putative transcription factor binding sites in the 5 kb region upstream of the human *RUNX2* gene promoter (NM_001015051.3) and identified eight potential STAT5 binding sites (**Supplemental Table 2**). We sought to determine whether STAT5 is bound to the *RUNX2* gene promoter and focused on the two most proximal predicted STAT5 sites located at −1371 bp and −193 bp upstream of the transcriptional start site (TSS), as both have high homology to the tetrameric consensus STAT5 binding site and show an elevated likelihood of being functional sites (**Figures 4A** and **4B**). As inflammation and inflammatory signaling pathways are known to contribute to CAVD pathogenesis^1,66^, and *TERT* expression was induced via that inflammatory transcription factor NF-κB (Supplemental Figure 2D), we used bacterial lipopolysaccharide (LPS) as a broad method to activate inflammatory response pathways in hVICs. We found that LPS was sufficient to induce calcification of hVICs and that TERT-positive cells accumulated near calcified nodules (**Figure 4C**). Protein analysis showed that OST and LPS significantly upregulate TERT, RUNX2, and STAT5 proteins (**Figure 4D**).

TERT protein levels were elevated in both the cytosol and nucleus of cells upon OST stimulation (**Supplemental Figure 5A**). Under NT conditions, STAT5 is observed in the membrane and cytosol but translocates and accumulates in the nucleus of hVICs in OST (**Supplemental Figure 5B**). Next, we used genetic and pharmacological approaches to determine the requirement of STAT5 for the calcification of hVICs. The small molecules StafiA-1 and StafiB-1 specifically inhibit STAT5A and STAT5B, respectively.^67,68^ We found that StafiA-1 significantly inhibited the calcification of control hVICs during a 21-day OST assay and that this inhibition was greater than the inhibition observed with StafiB-1 (**Figure 4E**). In line with these findings, we also observed that inhibition of STAT5A significantly reduced the expression of *RUNX2* at d7 of osteogenic differentiation of hVICs, suggesting that STAT5A is a main participant during the osteogenic reprogramming of hVICs (**Figure 4F**). Similarly, StafiA-1 significantly reduced calcification in hCASMCs and hMSCs (**Supplemental Figures 5C** and **5D**). Lentiviral-mediated delivery of shControl or shSTAT5 constructs showed that depletion of STAT5 significantly attenuated calcification and *RUNX2* expression (**Figure 4G** and **4H**), confirming an essential role for STAT5 in the osteogenic reprogramming of hVICs. Similar to TERT (Supplemental Figure 4D), knocking down STAT5 after 7 or 14 days of OST treatment did not reduce calcification (**Figure 4I**), indicating that STAT5 activity is required during the initial steps of osteogenic reprogramming. Others have established that TERT can interact with transcription factors to regulate gene expression.^17,20,62,63^ We assessed whether TERT and STAT5 directly interact. Three methods were utilized: proximity ligation assay (PLA), co-immunoprecipitation (IP), and exogenously expressing TERT and STAT5 reporters. PLA can detect protein-protein proximity <40 nm apart.^69^ PLA revealed that OST significantly increased TERT and STAT5 colocalization in control hVICs after 21 days of OST treatment relative to NT condition (**Figure 4J**). Focusing on nuclear co-localization, TERT/STAT5 puncta are significantly elevated in the nucleus of calcifying hVICs relative to the NT condition (**Supplementary Figure 5E**). Protein IP shows that TERT and STAT5 physically interact in hVICs upon OST treatment relative to NT condition (**Figure 4K**). A Dual-Luciferase reporter assay shows that the human *RUNX2* promoter, containing the −194 and −1371 putative STAT5 binding sites, is upregulated early (d3) under OST treatment relative to NT, and that this upregulation is abrogated with pharmacological inhibition of STAT5A, validating the functionality of the predicted STAT5 binding in the *RUNX2* promoter (**Supplementary Figure 5F**).

**Figure 5.**
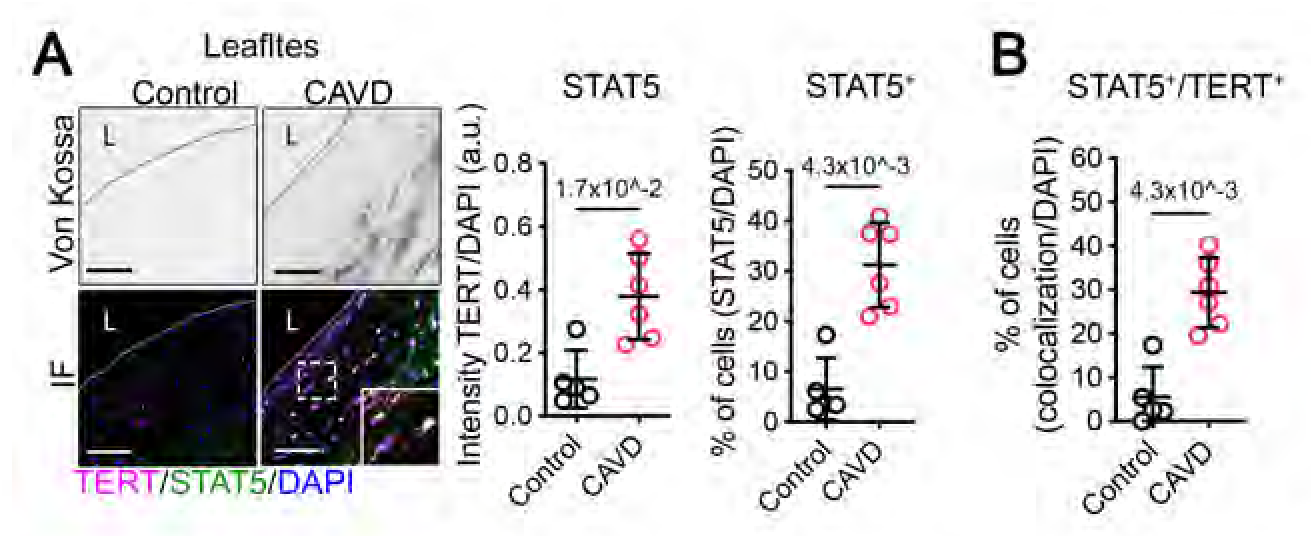
TERT and STAT5 are upregulated and colocalized in CAVD tissue. (A) Representative serial sections of Von Kossa (dark precipitation) and TERT and STAT5 immunofluorescent staining in control and CAVD tissues. Scalebar 100 μm. STAT5 intensity and STAT5-positive cells (STAT5+) are quantified on the right panels. (B) Quantification of STAT5-TERT-positive cells. Data are shown as means ± SD, n = 5 control, n = 6 CAVD. P values were calculated with the Mann-Whitney U test and indicated on each graph. L = lumen.

Furthermore, an electrophoretic mobility shift assay (EMSA) confirmed the physical binding of STAT5A to DNA oligos containing the predicted −194 STAT5A binding site on the RUNX2 promoter (wt RUNX2 oligo). This interaction was validated by challenging the oligo-STAT5 complex with increasing amounts of unlabeled (cold) oligos. Moreover, the binding was entirely abrogated by incubating the STAT5 extract with oligos where the STAT5A binding site was entirely randomized (mut RUNX2 oligo) (**Supplementary Figure 5G**). Supporting a role for STAT5 in osteogenic reprogramming, STAT5 protein levels were significantly lower in the aorta of *Tert^-/-^* mice fed the A6P6 diet compared to the aorta from *Tert^+/+^* mice (**Supplemental Figures 5H**), and these findings were replicated in the ex vivo calcification assay (**Supplemental Figure 5I**). These experiments demonstrate that STAT5 directly acts on DNA in the *RUNX2* promoter and contributes to osteogenic reprogramming in cardiovascular tissues.

While the domains of TERT were initially defined by their role in the telomere-extending functions of TERT, protein partners required for the known non-canonical functions of TERT have been found in several domains throughout the protein.^70^ Further, sites for protein binding partners to STAT5, such as epigenetic modifiers and transcriptional regulators, have been found throughout the protein.^71^ We used AI-assisted technology to generate tertiary structures of TERT and STAT5 dimers and predict how they may interact. AlphaFold^72^ mapped the interaction between human TERT and dimerized human STAT5A, and these binding data were visualized using HDOCK^73^ (**Supplemental Figure 5J** and **5K**). To gain a more comprehensive and detailed understanding of the TERT/STAT5 interaction we will utilize a next-generation fluorescence-based complementation system developed by Tebo and Gautier in 2019 (splitFAST).^74^ This system has very low background fluorescence and can be temporally regulated, enabling the visualization of transient TERT/STAT5 interactions in living cells or fixed cells, to provide a scalable model that can be used to screen for compounds that block the TERT/STAT5 interaction. We fused the human cDNA of TERT and STAT5 to NFAST and CFAST splitFAST reporters, respectively, to generate TERT-NFAST and STAT5A-CFAST fusion proteins. This approach allows monitoring of the formation and dissociation of a protein assembly.^74^ Control hVICs were co-transfected with TERT-NFAST and STAT5A-CFAST plasmids and 5 μM HBR-3,5DOM (4-hydroxy-3,5-dimethoxy benzylidene rhodamine) administered at d3, d5, and d7 of OST or NT. We found that the TERT/STAT5 splitFAST signal was highly elevated in calcifying control hVICs relative to the NT condition as early as d3 throughout d7, supporting the early formation of the TERT/STAT5 complex in calcifying hVICs (**Figure 4L**). Together, these approaches confirm TERT and STAT5 interaction and translocation into the nucleus during osteogenic reprogramming.

To assess the function of TERT-STAT5 interaction, we performed chromatin immunoprecipitation analysis (Ch-IP). We found that STAT5 is significantly enriched on both putative STAT5 binding sites in the *RUNX2* promoter during osteogenic reprogramming (**Figure 4M**). *RUNX2* transcription is initiated from two distinct promoters: the distal promoter region P1, located upstream of exon 1, and the proximal P2 promoter, located upstream of exon 2.^75,76^ The P1 promoter is highly active and required for osteoblast progenitor expansion, osteoblast maturation, and bone formation, while the P2 promoter-derived isoform is expressed in osteoblastic and non-osteoblastic mesenchymal cells.^77–79^ Genome-wide CUT&Run analysis of the histone marker H3K4me3 showed that RUNX2 promoters P1 and P2 are significantly upregulated during osteogenic differentiation on human VICs, while Gene Ontology (GO) pathway analysis revealed that genes related to the telomerase complex, chromatin remodeling, and metabolic pathways are upregulated during osteogenic differentiation of hVICs (**Supplementary Figures 5L and 5M**, GEO accession GSE280657). We assessed STAT5 binding to the *Runx2* P1 and P2 promoters during osteogenic differentiation using CUT&Tag analysis in the *Tert^+/+^* and *Tert^-/-^* mVICs. We found that STAT5 binds to the *Runx2* P1 and P2 promoters during osteogenic differentiation and that STAT5 was absent from these regions in *Tert^-/-^* mVICs, highlighting the Tert requirement for Stat5 binding to *Runx2* gene promoters (**Figure 4N**, GEO accession GSE274226). By comparing *Tert^+/+^* and *Tert^-/-^* mVICs CUT&Tag sequencing, we found that Stat5 occupies promoter, intragenic, and intergenic regions throughout the genome (**Figure 4O**). GO pathway analysis revealed that genes with high Stat5 binding participate in biological processes related to protein phosphorylation, JAK/STAT signaling, inflammation, osteoblast differentiation, SMC function, and chromatin remodeling (**Figure 4P**). Together, these data identify that TERT’s effects are mediated via STAT5 binding on the *Runx2* gene promoter and the promoters involved in several pathways known to be operative during osteogenic reprogramming of cardiovascular cells.

### TERT and STAT5 are upregulated in calcified tissue

Confirming our in vitro, ex vivo, and in vivo data above, in human CAVD tissue we found that STAT5 protein levels are significantly elevated and localized to areas of calcification in the calcified leaflet alongside TERT distribution. Additionally, the number of STAT5-positive cells was significantly higher in CAVD tissue compared to control tissue (**Figure 5A**). Furthermore, the number of TERT/STAT5 co-positive nuclei was significantly greater in CAVD valves compared to control valves (**Figure 5B**). These data support our in vitro investigations and indicate that TERT/STAT5 interaction contributes to human CAVD pathogenesis.

### TERT interacts with SMARCA4 during the osteogenic reprogramming of VICs

Studies by others have shown that TERT facilitates chromatin accessibility and transcription by recruiting SMARCA4 (previously referred to as BRG1), the catalytic subunit of the SWI/SNF chromatin remodeling complex, to specific DNA loci.^18,80,81^ We investigated whether SMARCA4 plays a role in the osteogenic reprogramming of hVICs via interacting with TERT. There was no difference in SMARCA4 protein staining in the aorta of *Tert^+/+^* and *Tert^-/-^* mice fed the A6P6 diet, nor in aortic rings ex vivo (**Supplemental Figures 6A** and **6B**). However, in hVICs, expression analysis revealed significant upregulation of *SMARCA4* transcripts as early as d5 of OST stimulation (**Supplementary Figure 6C**). Immunoprecipitation showed that TERT and SMARCA4 physically interact in hVICs upon OST treatment (**Supplementary Figure 6D**). Immunofluorescence staining showed that TERT and SMARCA4 are detected and colocalized in cells in the calcified human leaflet (**Supplementary Figure 6E**), mirroring TERT and STAT5 distribution in human CAVD tissue. These findings suggest that SMARCA4 plays an active role in the osteogenic reprogramming of VICs.

**Figure 6.**
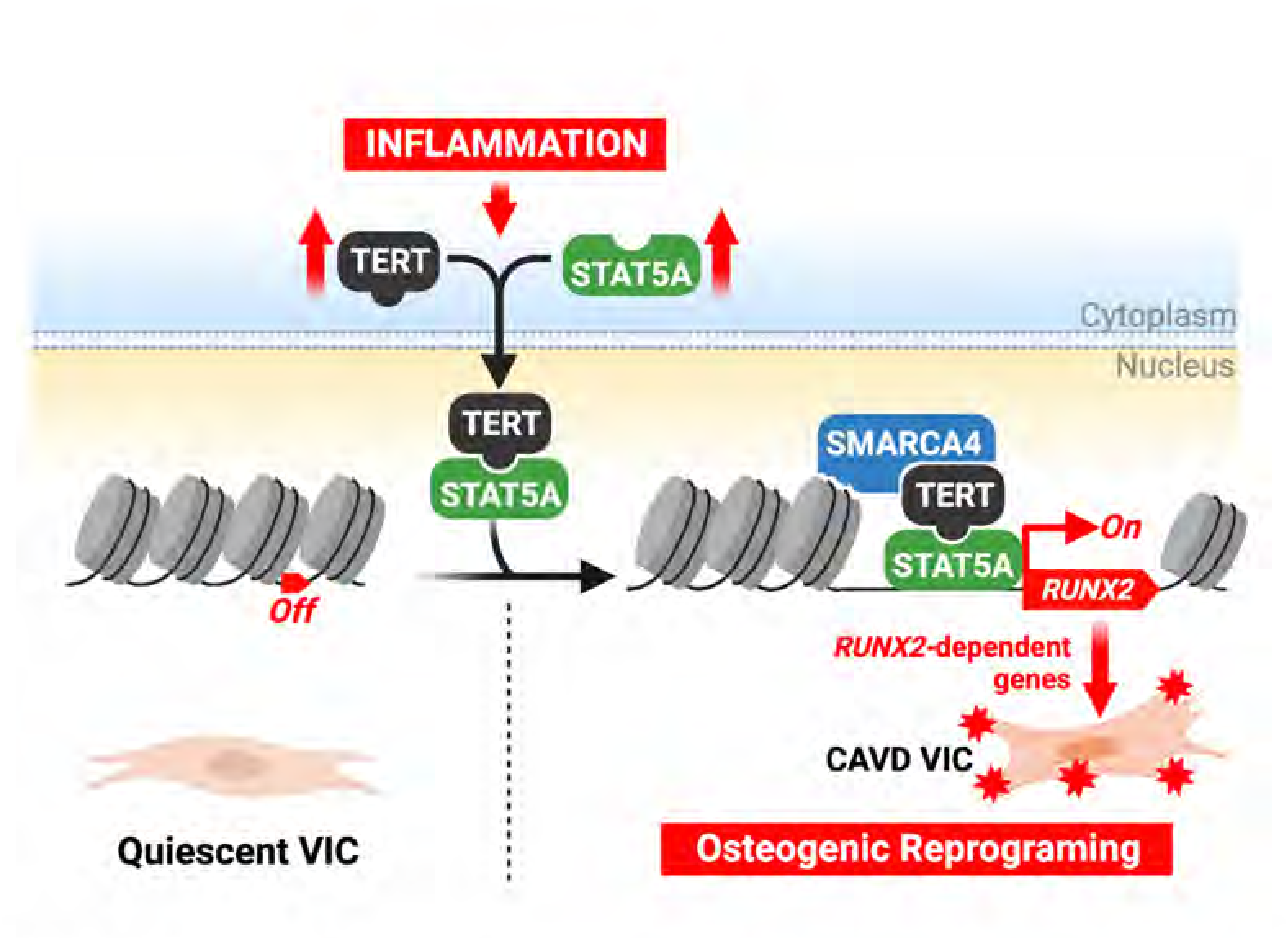
TERT/STAT5 promotes osteogenic reprogramming. Inflammatory signals promote the upregulation and interaction of TERT and STAT5A. The TERT/STAT5 complex translocates into the nucleus, where TERT interacts with SMARCA4 (also known as BRG1), which facilitates a permissive chromatin configuration, and STAT5A binds specific sites on the RUNX2 promoter, leading to the osteogenic reprogramming of valve interstitial cells during the early stages of CAVD pathogenesis.

## DISCUSSION

This study investigated the early steps required for the osteogenic reprogramming of human valve interstitial cells in CAVD and establishes a critical role for TERT and STAT5 in this process (**Figure 6**). Specifically, our data show that 1) TERT is upregulated in CAVD tissue and elevated in primary VIC lines stimulated with osteogenic media. While TERT is elevated in these calcified tissues and cells, telomere length is not impacted, nor is there evidence of cell senescence; 2) TERT is required for the calcification of VICs, CASMCs, and MSCs. The requirement for TERT occurs early in the process of osteogenic reprogramming, as knocking down TERT protein levels after seven days of osteogenic stimulation did not deter calcification. These osteogenic reprogramming activities of TERT are independent of its telomerase enzymatic activity, as *Terc^-/-^* cells or the use of the TERT reverse transcriptase inhibitor did not deter calcification. Further, TERT co-immunoprecipitated with the chromatin remodeling enzyme SMARCA4, and these two proteins were also found in the nucleus of CAVD tissues; 3) Inflammatory signals are known to contribute to CAVD pathogenesis, and we found such signaling can increase TERT levels, as well as the transcription factor STAT5. In silico modeling predicted TERT/STAT5 interaction, which was proven experimentally with immunoprecipitation, proximity ligation assay, and a fluorescence complementation system developed for the visualization of transient protein-protein interactions, and like TERT, genetic knockdown of STAT5, or specific inhibition of STAT5 activity, prevented calcification. We found that STAT5 directly binds the DNA of the RUNX2 promoter, as well as genes in pathways related to chromatin remodeling and cellular proliferation and differentiation. Lastly, the requirement for TERT was established in in vivo and ex vivo calcification models. Collectively, these experiments support a novel role for TERT and STAT5 in driving CAVD pathogenesis and cardiovascular reprogramming and identify a unique, druggable target for cardiovascular calcification.

TERT is well-known for its canonical telomere-extending activity, yet TERT also regulates gene expression and chromatin remodeling independent of affecting telomere integrity. TERT interacts physically with several transcription factors to promote the expression of several genes, including β-catenin, to stimulate genes involved in bone development.^15,16,62^ In a wire-injury restenosis model, TERT binds to the transcription factors SP1 at the *VEGF* promoter and E2F1 at *MCM7*, *CCNA1*, and *PCNA* promoters, stimulating SMC proliferation and neointima formation.^20,56,63,82^ We did not detect proliferative or migratory differences between control and CAVD VICs despite elevated levels of TERT at baseline (Figures 2B and 2C), indicating cell-specific and stress-specific functions of TERT.

In addition to supporting transcription factor function, TERT can act as a cofactor to orchestrate chromatin remodeling machinery. Studies have shown that TERT serves as a chaperone to facilitate the recruitment of SMARCA4 and histone acetyltransferases to stimulate chromatin accessibility and transcription or to maintain heterochromatin structure,^18,80,81^ and loss of TERT abrogates the cellular response to DNA double-strand breaks and alters the overall chromatin configuration without affecting telomere integrity.^17^ We now show that TERT is upregulated in calcified valve tissues and VICs under osteogenic and inflammatory stimuli. TERT interacts with STAT5, which binds to the *RUNX2* gene promoter and drives osteogenic reprogramming of the aortic valve cells. TERT depletion reduced *RUNX2* expression and calcification in several cell types, and TERT, STAT5, and the chromatin remodeling enzyme SMARCA4 co-immunoprecipitated in OST-treated VICs (Supplementary Figure 6). Further, TERT and SMARCA4 co-localized in the nucleus of CAVD tissues. These discoveries, along with the findings of others showing that *TERT* overexpression drives osteogenic differentiation in human MSCs,^21,22^ support a broad role for TERT serving as a co-factor bridging chromatin remodeling complexes and transcription factors to facilitate euchromatin formation to modulating cellular phenotypic transition and osteogenic reprogramming.

Telomere length is considered a biomarker of aging, yet it varies widely between people of the same age, even twins.^23^ Several studies have utilized circulating leukocytes to investigate whether shortened telomeres correlate with various vascular disease states.^24,25^ One study correlated telomere length in circulating leukocytes with aortic stenosis and found that leukocyte telomere length was slightly shorter in individuals with aortic stenosis than in age-matched controls.^26^ However, telomere length in leukocytes does not indicate global telomere length in an individual, telomerase complex activity, or TERT protein function in all cell types. Further, somatic mosaicism and clonal hematopoiesis discoveries underscore the importance of not ascribing telomere length in one cell type to telomere length and/or TERT activities in another.^28^ Short telomeres in circulating leukocytes may reflect increased cell turnover due to systemic inflammation, as leukocytes are highly proliferative, and telomere attrition is highly likely to be a tissue- and disease-dependent process. In this regard, Huzen et al. compared the telomere length of the leukocytes of age-matched controls and patients with atherosclerosis and the telomere length of the cells in the atherosclerotic plaques of the patients.^27^ They found that while telomere length in leukocytes from the atherosclerosis patient group was significantly shorter than in the control group, telomere length in the atherosclerotic plaques was significantly longer, indicative of increased telomerase activity. Thus, total leukocyte telomere length does not represent telomere length in the diseased tissue. Our study investigated TERT protein levels directly in the valve tissue and patient-specific VICs. We determined that the expression levels of TERT and osteogenic markers in CAVD tissue and cells were elevated, without changes in telomere length, proliferation, and DNA damage markers (Figures 1-2, Supplemental Figures 1-4), suggesting non-canonical TERT activity in these cells.

VIC calcification shares aspects of the transcriptional program observed during the differentiation of MSCs into osteoblasts, including the upregulation of *RUNX2, BGLAP, TNAP*, and the secretion of bone-forming proteins and accumulation of calcium minerals.^10,12,83^ Likewise, signals such as mechanical stress, disturbed flow, and inflammation promote VIC calcification and the osteogenic differentiation of MSCs.^84^ These observations suggest that VICs switch from their quiescent state and undergo an osteogenic reprogramming akin to the differentiation of MSCs into osteoblasts, acquiring a calcifying phenotype. The mechanisms coordinating the early molecular events in these cell transitions have yet to be fully understood. We have hypothesized that fully differentiated cells can de-differentiate into a transitory, multipotent stem cell-like state and then acquire an osteogenic phenotype.^85^ VICs exhibit the potential to differentiate into osteogenic, chondrogenic, or adipogenic lineages, properties comparable to mesenchymal stem cells.^86^ We chose to investigate TERT in the osteogenic reprogramming of VICs as TERT is highly expressed in stem cells, and its overexpression primes mesenchymal stem cells into the osteogenic lineage.^21,22^ Our data supports the role of TERT in the osteogenic reprogramming of VICs, whereby TERT serves as a bridge between DNA-binding transcription factors and chromatin remodeling enzymes.

The osteogenic differentiation of stem cells is orchestrated by RUNX2, which is also required for the osteogenic reprogramming of valvular and vascular cells.^40,41^ In a recent transcriptomic study, Xu et al. identified 14 cell populations in aortic valve tissue samples from two control and four CAVD valves.^87^ The severity of CAVD was assessed via ultrasound imaging, which detects leaflet mobility; however, examining the macroscopic pictures and histological staining of valve leaflets used in this analysis shows that while these valves are indeed stiff, they do not exhibit robust nodules of calcification as do the valves in our data set (Figures 1, 5, and Supplemental Figure 1). Analysis by Xu et al identified three VIC populations defined by expression of *FOS, HSPA6, and SPARC.* However, cells from six valves were pooled, with 10.4% of cells from control tissue and the remaining 89.6% from CAVD tissues. Bulk transcriptomic data from these tissues show only a small 1.49-fold increase in *RUNX2* gene expression in CAVD tissues, in stark contrast with our more stringent analysis of *RUNX2* expression by quantitative PCR (Figure 2, Supplemental Figure 4).^10–12^ Inspection of the publicly available bulk RNA-sequencing data (PRJNA552159) showed no increase in *TERT* transcript expression. While the hemodynamic data showed those valves had undergone fibrotic remodeling, few calcifying cells were present, as seen by their histological staining. This lack of calcification may explain why TERT expression was undetected in their study. In the general population, aortic sclerosis commonly goes undetected until a patient develops symptoms such as dyspnea, angina, and syncope, as the stiffness and remodeling of the leaflet obstructs ventricular outflow. A population-based study assessed aortic valve disease in 552 randomly selected men and women aged 55 to 86. Using imaging and Doppler echocardiography, mild calcification was observed in 40% of the individuals, severe calcification in 13%, and symptomatic stenosis in only 2.2%.^88^ This population-based study illustrates how common aortic valve remodeling is and highlights the importance of using multiple modalities to accurately determine valve pathology (i.e., fibrotic vs calcified). In our study, a portion of each patient valve was used for cell isolation while the rest was utilized for calcification assessment by histological staining, following a workflow that we developed that guarantees that cells isolated from surgically removed and postmortem tissues retain their proliferative capacity and endothelial and interstitial phenotypes in culture.^37^ While unbiased sequencing approaches may help uncover novel mechanisms contributing to disease states, they are skewed towards genes exhibiting the largest changes in gene expression in a snapshot of time and can miss short-lived transitory events.

The function of STAT5 in bone formation is multifaceted. While it was shown that STAT5 could promote the differentiation of MSCs into osteoblasts by upregulating the osteogenic genes in a Jak2-dependent manner, a *Stat5a* global knockout mouse exhibited elevated bone mass and mineral density.^89,90^ These conflicting reports indicate that the role of STAT5 in bone formation and remodeling is complex, and more studies are required. STAT5 has diverse functions as a transcription factor: it interacts with histone acetyltransferases and transcription factors such as the glucocorticoid receptor, SP1, YY1, and C/EBPβ to stimulate gene expression,^91–93^ and studies in cancer cells identified that STAT5 can induce *TERT* expression.^94,95^ Using in silico modeling we found multiple putative STAT5 binding sites in the *RUNX2* P1 promoter region, which were confirmed with Ch-IP assays. This supports our hypothesis that STAT5/TERT acts to initiate the osteogenic reprogramming of VICs, establishing a novel mechanism in CAVD pathogenesis whereby TERT/STAT5 bind to each other and activate osteogenic gene transcription, which drives the early events in cellular osteogenic reprogramming. Cell phenotype reprogramming requires chromatin restructuring to allow for the transcription of repressed genes. As observed in other cell types, we found TERT co-immunoprecipitated with SMARCA4, as well as STAT5, and genetic depletion of TERT or STAT5 midway through the OST assay (i.e., after d7) did not perturb the calcification of cells. Our observations strongly supports that TERT complexes with SMARCA4 to remodel chromatin and with the STAT5 transcription factor to bind and activate *RUNX2* gene transcription. We explored the potential physical direct interaction between TERT and STAT5, and we propose that the RNA binding domain of TERT is the most likely candidate to interact with STAT5 (Supplemental Figure 5). This location is of interest, as we found that *Terc^-/-^ mVICs* robustly calcify; this begs the question of whether the loss of *lncTERC* binding is what enables or regulates the non-canonical activities of TERT. Studies identifying critical domains for interaction are currently ongoing.

The strengths of this study lie in the utilization of patient-specific tissues and cell lines generated from valve leaflets, and the use mouse models and knockout cells for mechanistic studies. While our results are robust, it is important to address the limitations of our study. Firstly, we worked with a limited number of human samples obtained from age- and sex-matched donors. Due to this limited sample size, we limited our statistical analyses to non-parametric analysis, comparing two to four groups, and no mixed models were included. Regardless, one strength of our study is the multiprong approach to determining the role of TERT and STAT5 in vascular calcification. Secondly, STAT5A and STAT5B are two different genes, with 96% of their amino acid sequence shared and their activities redundant.^32^ We have determined that VICs express both variants; however, our antibodies and shRNA constructs could not differentiate between the two variants. Thus, the two isoforms may interact with TERT to bind the *RUNX2* gene promoter. STAT5 is activated by multiple cytokines, including IL-2, IL-6, and TNF-α, and it was shown that IL-2 specifically activates STAT5 to induce TERT expression in cancer cells.^94,96^ Whether STAT5 first induces TERT expression and then TERT and STAT5 cooperate to control the expression of *RUNX2* needs to be determined. Thirdly, there is redundancy regarding STAT1/3/5 functions and signaling.^97^ Our study was not designed to address all the complexities of STAT signaling per se but rather to investigate the role of TERT as a key protein that orchestrates the osteogenic reprogramming of aortic valve cells; our future studies will explore the role of STAT5 further.

Our work highlights the importance of primary human tissue-based studies. Murine models are not ideal for studying non-genetic drivers of CAVD, as mice do not develop CAVD de novo.^98,99^ Mouse leaflets’ structure differs from human leaflets as they lack a trilaminar organization and present a melanocyte cell population not present in human valves.^100–102^ Recent elegant work from the Giachelli group showed that in mice, calcification happens in the sinus wall and, to a lesser extent, at the leaflet hinge, the region between the leaflet VICs and SMCs of the aorta, with no detectable calcification in the aortic valve leaflets after 26 weeks of a high-fat diet in a *Ldlr* knockout mice model.^103^ This distribution of aortic root calcification is also observed in a mouse model described by a Honda et al which detected calcification in the sinus wall 16 weeks post wire-induced mechanical injury.^98^ Due to the lack of aortic valve leaflet calcification in atherosclerotic or mechanical injury murine models, we utilized a adenine/phosphate diet-induced model of medial arterial calcification (MAC).^58^ MAC is not associated with lipid deposition, fibrous cap formation, or intimal hyperplasia; instead, MAC stems from the progressive accumulation of calcium and phosphate within the arterial walls.^104^ These structural features distinguish MAC from atherosclerosis and are more representative of the pathogenesis of valve leaflet calcification.^2,104^

This study has uncovered the early molecular events that enable the osteogenic reprogramming of the interstitial cells of the aortic valve. Current CAVD therapies are surgical, limited to mechanical or bioprosthetic valve replacement, and performed only when the disease has progressed to the advanced point of affecting blood flow and heart function. Our data show that perturbing TERT or STAT5 can inhibit calcification; however, broadly inhibiting one or the other of these proteins may have deleterious off-target effects. Understanding the specific binding domains and other protein binding partners of TERT and STAT5 will enable the development of small molecules to block their interaction, which is a more feasible therapeutic strategy for aortic valve calcification than global inhibition. CAVD affects over 9.5 million people in the U.S. and has an associated 56-67% 5-year mortality rate; hence, the need for early, non-surgical therapies is great.^4^ This study uncovers a novel and targetable pathway that may be leveraged to halt or prevent CAVD progression.

## ACKNOWLEDGEMENTS

We want to acknowledge and thank the Center for Organ Research and Education for their help and support, as well as tissue donors and their families for making this study possible. We thank Dr. Alison Tebo for her guidance and assistance in constructing the SplitFAST plasmids tool used in this study. We would also like to thank Kristin Konopka, tissue bank coordinator at The Department of Cardiothoracic Surgery, UPMC. We thank Jason Dobbins for his critical reading of this manuscript.

## SUPPLEMENTAL MATERIAL

## METHODS

### Availability of Materials

We abide by the NIH Grants Policy on Sharing of Unique Research Resources, including the NIH Policy on Sharing of Model Organisms for Biomedical Research (2004), NIH Grants Policy Statement (2003), and Sharing of Biomedical Research Resources: Principles and Guidelines for Recipients of NIH Grants and Contracts (1999), and the Bayh-Dole Act and the Technology Transfer Commercialization Act of 2000. De-identified human cell lines and tissues generated in our laboratory will be made available for non-commercial research per established University of Pittsburgh Office of Research IRB and MTA protocols.

### Ethics

Human aortic valves were collected from subjects enrolled in studies approved by the institutional review board (IRB) of the University of Pittsburgh per the Declaration of Helsinki. No bicuspid valves were used in this study. Mice used for the generation of cell lines were given veterinary care by the University of Pittsburgh Division of Laboratory Animal Resources, which adheres to the NIH policy on the Animal Welfare Act and all other applicable laws. Facilities are under the full-time supervision of veterinarians and are AAALAC-accredited. Our protocols follow the AVMA Guidelines on Euthanasia. All animal use was approved by the Institutional Animal Care and Use Committee at the University of Pittsburgh.

### Human Tissue Collection and Isolation

Surgical specimens from humans were collected from subjects who consented and enrolled in studies approved by the institutional review board (IRB) of the University of Pittsburgh per the Declaration of Helsinki. Personnel involved with specimen handling underwent extensive institutional training. Cadaveric donor tissues were obtained via the Center for Organ Recovery and Education (CORE) and were approved by the University of Pittsburgh Committee for Oversight of Research and Clinical Training Involving Decedents (CORID). A detailed protocol describing tissue collection hand handling was published previously.^1^ Briefly, valves were collected from valve replacement surgeries or recovered from cadaveric organs and stored in cold Belzer UW Cold Storage Transplant Solution (Bridge to Life) at 4°C for transporting. Aortic roots were excised and washed with a sterile rinsing solution (sterile PBS supplemented with 2.5 μg/mL of fungicide, 0.05 mg/mL of gentamicin, and 5 μg/mL of bactericide). Leaflets were unbiasedly selected for VIC isolation, Von Kossa staining for calcification, and snap freezing for RNA collection. Tissues were processed as close as possible to the extraction time to guarantee the best yield of cell recovery.

### Human Cell Isolation

We established patient-specific lines by using the same valves for histopathology, RNA, and cell isolation. Human primary aortic valve interstitial cells (VICs) lines were generated from male and female subjects as previously described.^1^ Briefly, leaflets were washed with PBS containing 10 mg/ml gentamicin (GIBCO) and 250 μg/ml fungizone (GIBCO) and dissociated with 0.1% collagenase II at 37 C and 5% of CO2 for 18 hours. Then, the tissue was further dissociated by gently mixing it by pipetting with a serological pipette to ensure the release of VICs and then passed through a 0.70 μm filter to remove debris. Cells were pelleted and then resuspended in Dulbecco’s Modified Eagle’s Medium (DMEM) supplemented with 10% fetal bovine serum (FBS) and 1X penicillin-streptomycin (P/S). Human coronary artery smooth muscle cells (CASMC) were obtained from the patient coronary as previously described.^2^ Briefly, vessels were washed with PBS containing 10 mg/ml gentamicin (GIBCO) and 250 μg/ml fungizone (GIBCO). Vessels were cut open to expose the lumen, and intima and adventitia were gently scrapped. Vessels were then sectioned and dissociated with 0.1% collagenase II for 3 hrs. Cells were pelleted and then seeded and expanded.

### Murine tissue and cell isolation

All murine cell lines are generated from a single animal. Males and females, 6 to 8 weeks old, *Mus musculus Terc^-/-^* (Jackson Labs, strain # 004132) mice were utilized for cell isolation. Males and female *Tert^-/-^* (Jackson Labs, strain # 005423, 10-12 weeks old) mice were bred with wild-type mice (Jackson Labs, 000664) to produce *Tert^+/-^* heterozygous mice. Heterozygous *Tert^+/-^* breeding regimes were used to prevent telomere attrition. PCR confirmed genotype. Mouse BM-MSCs were isolated from femurs, and tibias were dissected from three-month-old *Tert^+/+^* or *Tert^-/-^* mice. The marrow was rinsed out of the bones with MSC media, and cells were plated and expanded as described previously.^3^ Mouse VICs were isolated from the hearts of three-month-old *Tert^+/+^ or Tert^-/-^* or mice and were removed, dissected, and valve leaflets removed. Cells were isolated and expanded as described previously.^4^ Animals were handled by designed members in our lab. Tissue and cell isolation were carried out by a different team to ensure randomization. Mice were given veterinary care by the University of Pittsburgh Division of Laboratory Animal Resources, which adheres to the NIH policy on the Animal Welfare Act and all other applicable laws. Facilities are under the full-time supervision of veterinarians and are AAALAC-accredited. Our protocols follow the AVMA Guidelines on Euthanasia. All animal breeding and isolations were approved by the Institutional Animal Care and Use Committee at the University of Pittsburgh. We made attempts to minimize the number of mice required to complete experiments.

### Cell Culture

VICs lines were expanded in Dulbecco’s Modified Eagle’s Medium supplemented with 10% fetal bovine serum and 1X penicillin-streptomycin. Cells were used between passages 4 and 15. Growth media was changed every three days, and cells were split 1:2 when confluent. Human coronary artery smooth muscle cells (CASMCs) lines were expanded in smooth muscle media SMGM (CC-3181) supplemented with BULLETKIT (CC-4149). Human mesenchymal stem cells (MSCs, PT-2501, Lonza) were expanded on Minimum Essential Medium alpha without nucleosides and supplemented with 10% fetal bovine serum and 1X penicillin-streptomycin and used between passages 4 and 10. Cells are routinely tested for mycoplasma contamination.

### Osteogenic Assay

For osteogenic experiments, 250,000 cells per 9.5 cm^2^ were seeded and treated with osteogenic media (OST, Gibco Minimum Essential Medium alpha with nucleosides, 10% FBS, 1X P/S, 10 mM glycerol phosphate, 50 μM ascorbic acid 2-phosphate, and 100 nM dexamethasone).^5^ No treatment media consisted of Gibco Minimum Essential Medium alpha with nucleosides, 10% FBS, 1X P/S. Media was replaced every four days and prepared fresh every time. BiBR1532 (2981, Tocris) was added to the media at 1, 10, or 100 nM every two days during the OST treatment.^6^ Inhibitors StafiA-1 and StafiB-1 (HY-136546, HY-112647 MedChemExpress) were used at 10 μM. Inhibitors BAY-117082 (catalog number S2913, SelleckChem) and 10058-F4 (catalog number S7153, SelleckChem) were used at a maximum of 10 and 50 10 μM, respectively. Inhibitors were added every two days for the duration of the OST treatment. For inflammatory assays, we used 2.5 μg/mL of LPS from Escherichia coli O111:B4 (Sigma-Aldrich) every two days for the duration of the OST treatment.

### SDS-Page and Immunoblotting

Cells were lysed in lysis buffer (1% CHAPS hydrate, 150 mM sodium chloride, 25 mM HEPES buffer), supplemented with 1x protease and phosphatase inhibitor cocktail (Sigma-Aldrich). Cells were scraped and transferred into microcentrifuge tubes, vortexed for 10 minutes, freeze/thawed for 5 cycles, then centrifuged at 12,000 x g for 10 minutes at 4°C, and supernatants were collected. Proteins were separated with TGX 4-20% stain-free polyacrylamide gel (Bio-Rad) in 1x Tris/Glycine/SDS buffer (Bio-Rad) and transferred to 0.2 um nitrocellulose (1620112, Bio-Rad) membrane in 1x Trans-Blot Turbo Transfer Buffer (Bio-Rad) using the Trans-Blot Turbo Transfer System (Bio-Rad) according to the manufacturer recommendations. The membranes were blocked in Odyssey blocking buffer (PBS, Li-COR) and immunoblotted overnight with primary antibodies against TERT (600-401-252, Rockland), STAT5 (9420S, Cell Signaling), RUNX2 (ab192256, Abcam), OPN (AF808, R&D systems), αSMA (ab5494, Abcam), TNAP (MAB2909, R&D systems), a-tubulin (926-42213, LI-COR), followed by secondary anti-rabbit or anti-mouse IgG antibody (926-68070, 926-68021, 926-32211, LI-COR). Primary and secondary antibodies were diluted in Odyssey blocking buffer with 0.1% Tween 20. Membranes were washed in PBS with 0.1% Tween 20. Immunofluorescence signals were detected with the Odyssey CLx system (LI-COR), and images were analyzed with Image Studio (Version 5.2, LI-COR).

### Immunofluorescent Staining on Fixed Cells

Cells were washed with PBS and then fixed with 4% paraformaldehyde (5025999, Electron Microscopy Sciences) for 15 minutes at room temperature, followed by 10 minutes of permeabilization with TRITON X-100 0.5% (Fisher Scientific). Cells were incubated at room temperature for 1 hour in a blocking buffer (PBS, 0.5% Triton-X 100, 5% FBS). Primary antibodies were diluted in the blocking solution and then incubated overnight at 4°C. Next, cells were washed three times for five minutes each with PBS containing 0.1% TWEEN. Fluorescent secondary antibodies were diluted in blocking solution and incubated for one hour protected from the light, then washed three times for five minutes each with PBS, 0.1% TWEEN 20 with a final wash in PBS. Specimens were finally mounted with Fluoroshield Mounting Medium with DAPI (Abcam) and imaged within 24 hours. F-Actin was stained with AlexaFluor488 Phalloidin (Molecular Probes) for 30 minutes, then washed with PBS before mounting. Calcium accumulation was determined with OsteoImage (Lonza) following the manufacturer’s recommendations. Images were obtained in an EVOS FL microscope (AMA3300, Life Technologies) at 20x with an EVOS Plan Fluor 20x/0.5 objective (AMEP 4698, Life Technologies) at 360/447 nm excitation/emission (DAPI), 470/525 nm excitation/emission (GFP), and 530/593 nm excitation/emission (RFP) filters. Representative images are selected based on quality, and if the assay includes quantification, the image that shows the average of the quantification across multiple images.

### Immunofluorescent Staining on Paraffin Sections

Human specimens were embedded in paraffin and cut to 10 μm thickness. Slides were warmed to 65°C, deparaffinized using xylene and graded alcohol baths, rehydrated in distilled H_2_O, and boiled for 20 minutes in antigen unmasking solution (H-3300, Vector Labs). Cooled samples were then washed with PBS for five minutes. Specimens were then blocked with PBS containing 3% fish skin gelatin and 10% horse serum for one hour. Primary antibodies were diluted in the blocking solution and incubated overnight at 4°C. Next, specimens were washed three times for five minutes each with PBS containing 3% fish skin gelatin and 0.1% TWEEN 20. Fluorescent secondary antibodies were diluted in a blocking solution, incubated for one hour, protected from the light, and washed thrice for five minutes each. Normal rabbit polyclonal IgG (Invitrogen, 31235) and Normal mouse IgG (Vector, I-200U) were used as a negative isotype antibody control at the same concentration of the specific antibodies. Specimens were finally mounted with Fluoroshield Mounting Medium with DAPI (Abcam) and imaged within 24 hours. Images were obtained in a EVOS FL microscope (AMA3300, Life Technologies) at 20x with a EVOS Plan Fluor 20x/0.5 objective (AMEP 4698, Life Technologies) at 360/447 nm excitation/emission (DAPI), 470/525 nm excitation/emission (GFP), and 530/593 nm excitation/emission (RFP) filters. Representative images are selected based on quality, and if the assay includes quantification, the image that shows the average of the quantification across multiple images.

### Alizarin Red Staining and Quantification

Cells were washed with PBS and then fixed with 4% paraformaldehyde (Electron Microscopy Sciences) for 15 minutes at room temperature, followed by two washes with deionized water. Fixed cells were then covered with 40 mM Alizarin Red S (Sigma-Aldrich) at pH 4.1 – 4.3 and gently rocked for 20 minutes at room temperature. Cells were then washed twice with deionized water to remove any unincorporated dye. After imaging, Alizarin Red S was extracted with 10% (v/v) acetic acid (Fisher Scientific) for 30 minutes, scraped into a microcentrifuge tube, vortexed, and then incubated at 85°C for 10 minutes. After chilling on ice for 5 minutes, the mixture was centrifuged at 20,000 x g for 15 minutes at 4°C. 500 μl of supernatant was transferred to a new tube, and 10% (v/v) ammonium hydroxide (Fisher Scientific) was then added to the supernatant. Absorbance was at 405 nm using a 96-well plate spectrophotometer.

### Von Kossa Staining

Human specimens embedded in paraffin were warmed to 65°C for one hour and then deparaffinized through xylene and graded alcohol incubations. Specimens were then rehydrated in distilled water. Specimens were stained using the Von Kossa Method of Calcium Kit (Polysciences, 24633-1) following the manufacturer’s instructions.

### Lentiviral Production and Cell Infection

Lentiviruses were produced by transfecting Dharmacon SMARTvector Lentiviral plasmids encoding shTERT TurboGFP (shTERT, V3SH11240-225610522). Turbo SMARTvector Non-targeting Control (shControl,VSC11707) into HEK293T cells following manufacturer recommendations. shSTAT5A (sc-29495-V) was obtained from Santa Cruz. The viral-containing supernatant was collected at 48h after transfection, filtered through a 0.45-μm filter, and stored at −80C. Human VICs, CASMCs, and MSCs lines were transduced with MOI of 5 in the presence of 0.8 μg ml^−1^ polybrene (Millipore) to enhance transduction efficiency.

### Transcriptional Analysis

Tissue RNA was isolated using Trizol (Life Technologies). Cell RNA was isolated using Quick-RNA MiniPrep (Zymo Research). RNA was treated with DNAse I (Zymo Research) in accordance with the manufacturer’s instructions. Reverse transcription was performed using MultiScribe Reverse Transcriptase system (43-112-35, Fisher). Sixteen ng of cDNA was used per reaction. qPCR was performed on a CFX Connect Real-Time System (Bio-Rad) using PowerUP SYBR Green Master Mix (A25741, Applied Biosystems) as follows: one cycle at 95 °C (10 minutes) and 40 cycles of 95 °C (20 seconds) and 58 °C (20 seconds) and 72°C (1 minutes). *GAPDH* or *18s* gene expression was used to normalize expression. Relative expression was calculated using the average threshold cycle number and the 2 ^(Ct(*housekeeping* gene)−Ct(*target gene*))^ formula. Primers are listed in Supplementary Table 3.

### Luciferase Assay

Human VICs (0.5×10^5^ cells/well) were seeded in 24-well plates and transfected with the RUNX2 Promoter GLuc-ON plasmid (HPRM32232-PG04, Genecopoeia) using Lipofectamine 3000 (L3000001, ThermoFisher). This plasmid encodes 1627 bp upstream of the human *RUNX2* gene NM_001015051. 48 hrs after transfection, the cells were treated with OST media for 72 hrs. Luciferase activities were measured using the Secrete-Pair Dual Luminescence Assay Kit (LF032, Genecopoeia), following to the manufacturer’s instructions. The results were expressed as relative light units (RLU) of luciferase to secreted alkaline phosphatase (SEAP)

### CUT&Run

The CUT&Run assay was performed using the CUTANA Kit V3 (14-1048, EpiCypher) according to the manufacturer’s instructions. Briefly, human VICs were cultured in osteogenic media for seven days. On the extraction day, 500,000 cells were harvested and tagged with concanavalin A-coated magnetic beads. The coated cells were incubated overnight with 0.5 μg of H3K4me3 antibody (Part number 13-0041k, EpiCypher) at 4°C. The following day, samples were incubated with pAG-MNase (Part number 15-1016, EpiCypher) for 2 hours at 4°C. DNA was purified and indexed with 14 PCR cycles to generate the libraries. Sequencing was performed on an Illumina NextSeq 2000, resulting in the generation of 60-bp paired-end reads. Raw data was processed and analyzed using CLC Genomic Workbench (version 24.0.1, Qiagen). Reads resulting from H3K4me3 binding were aligned to the human genome assembly GRCh38. Peaks were annotated to the nearest gene, and the binding between no treatment and osteogenic conditions was compared to identify osteogenic differentially regulated genes with a P-value of less than 0.05 (GEO accession number GSE274226). Visualization was conducted using the CLC Genomic Workbench. GO pathway analyses were performed with DAVID version 2023q4.^7,8^

### CUT&Tag

CUT&Tag assay was performed using the CUT&Tag-IT Assay Kit (Catalog No. 53160, Active Motif) following the manufacturer’s instructions. Briefly, wild-type *Tert^+/+^* and *Tert^-/-^* mouse VICs were cultured in osteogenic media for seven days. On the extraction day, 500,000 cells were harvested and tagged with concavalin-A-coated magnetic beads. Coated cells were incubated overnight with 1 μg of antibody against STAT5 (9420S, Cell Signaling) in 50 μl of total volume at 4°C with slow rotation. After washing, cells were incubated with guinea pig anti-rabbit antibody (Part No. 105465, Active Motif) at room temperature for 1 hour, followed by incubation with pre-assembled pA-Tn5 Transposomes at room temp for 1 hour (Part No. 105458, Active Motif). Tagmentated DNA was purified and indexed with 14 PCR cycles with Q5 Polymerase (part no. 104483, Active Motif) to generate the libraries. Sequencing was performed on an Illumina NextSeq 2000. 60-bp paired-end reads were generated. Raw data was processed and analyzed with CLC Genomic Workbench (version 24.0.1, Qiagen) supplied by the University of Pittsburgh. Reads resulting from STAT5 binding were aligned to the mouse reference genome mm10 assembly version GRCm39. The peaks were annotated to the nearest gene, and binding between no treatment and osteogenic conditions was compared to define osteogenic differentially regulated genes of those with a P-value less than 0.1 (GEO accession number GSE274226). Visualization was performed with IGV and CLC Genomic Workbench. GO pathway analyses were performed with DAVID version 2023q4.^7,8^

### Telomere Length Analysis

Tissue and VICs genomic DNA was isolated from passage 1 using DirectAmp Tissue Genomic DNA Amplification Kit (Denville Scientific). Telomere length was analyzed using real-time PCR as previously described.^9^ Briefly, genomic DNA was isolated following standard protocol, and 10 ng of gDNA per reaction was utilized. Samples were run in triplicate with 35 ng of DNA per reaction, and telomere repeats were amplified using PowerUP SYBR Green Master Mix (Thermofisher Scientific) on a CFX Connect Real-Time PCR System (Bio-Rad). Repeated amplification data were normalized to *RPLP0*/*36b4* as a single copy-gene. Primers are listed in Supplementary Table 3.

### Electrophoretic Mobility Shift Assay (EMSA)

HEK293 overexpressing STAT5 were lysed in ice-cold hypertonic non-denaturing buffer (20 mM Tris-HCl, pH 8.0, 137 mM NaCl, 1% Nonident P-40, and 2 mM EDTA) and microfuged at 14,000 rpm for 5 minutes at 4°C. The supernatant containing STAT5 was collected, and 6.8 μg of the extract was incubated for 30 minutes at 37°C with 20 fmol of biotinylated–end-labeled 60-mer double-stranded RUNX2 promoter containing the STAT5 centered at −194 5’-AAGCTATATCCTTCTGG<u>ATGCCAGGAAAGGCCTTACCACAAG</u>CCTTTTGTGAGAGAAAGA-3’ (underlined) binding site. The specificity of STAT5 binding to the wild-type probe was determined with mutated STAT5 binding site, 5′-AAGCTATATCCTTCTGGA<u>AACAACTGCTCGGCGAGACACAGT</u>CCTTTTGTGAGAGAAAGA-3’ oligonucleotide. Binding reactions were carried out with LightShift® Chemiluminescent EMSA Kit (ThermoFisher) according to the manufacturer’s instructions. Briefly, a final volume of 20 μL containing the STAT5 extract and the probe plus 1 μg poly(dI-dC), 100 mM MgCl2, 50% glycerol, and 1x propietary binding buffer. Bound complexes were separated on 6% acrylamide gel with TBE running buffer, crosslinked to a nylon membrane at 120mJ/cm^2^, and then visualized on BioChem Docs (Bio-RAD). No STAT5 binding was displaced by adding unlabeled (cold) wild-type oligonucleotide to the reaction. No STAT5 binding to the mutated probe was detected in our experiments, confirming the specificity of interaction between STAT5 and wild-type oligonucleotide.

### Proliferation Assays

Cell proliferation was evaluated using Trypan Blue incorporation in an automatized cell counter Countess II FL (Invitrogen). Briefly, cells were grown on alpha-MEM supplemented with 10% of FBS and penicillin and streptomycin cocktail (GIBCO) for the duration of the assay. Growth was quantified twice a week. Cell number during OST treatment was determined at the beginning and end of the assay.

### Senescence-Associated β-Galactosidase Assays

Cellular senescence was evaluated by senescence-associated β-Galactosidase (SA-β-Gal) activity assay. Briefly, cells were washed with PBS and then fixed with 4% paraformaldehyde (Electron Microscopy Sciences) for 5 minutes at room temperature and then washed twice with PBS. Next, cells were incubated in X-Gal Solution Mix containing 1 mg/mL of 5-Bromo-4-Chloro-3-Indolyl-Alpha-D-Galactopyranoside (American Bioanalytical), 40 mM Sodium Phosphate monobasic solution (Sigma-Aldrich), 40 mM Sodium Phosphate Dibasic solution (Sigma-Aldrich), 40 mM Citric Acid (Sigma-Aldrich), 5 mM Potassium Ferrocyanaide (Sigma-Aldrich), 5 mM Potassium Ferricyanaide (Sigma-Aldrich), 150 mM NaCl_2_, and 2 mM MgCl_2_, pH 6.0, for 16 hours at 37°C without CO_2_ to develop blue color. Then, cells were washed twice with PBS, and SA-*β*-gal positive cells were imaged and calculated based on randomly selected bright fields areas containing at least 200 cells per field as revelated by subsequent DAPI staining.

### Cell Migration Assays

VICs were seeded in a 12-well plate at a concentration of 1×10^5^ cells per well and were left until they reached 90% of confluence. The well surface was scratched with a 200 μl sterile pipette tip and washed with PBS to remove detached cells. Horizontal lines were drawn on the bottom outside of the well and used as a reference for alignment to obtain the same field for each image acquisition run. VICs media was added, and images were collected at different time points with a phase-contrast microscope using the horizontal lines as reference marks. Scratch area and scratch width were determined with the Wound Healing Size Macro Tool using ImageJ.^10^ Linear regression was used to compare the changes in the area and the average length of the scratch

### Chromatin Immunoprecipitation

Chromatin Immunoprecipitation (ChIP) was performed on cultured cells as previously described.^11^ Briefly, VICs were stimulated with OST for 14 days and then collected in 20 mM Na-butyrate dissolved in PBS, and cross-linking of DNA and proteins was performed with formaldehyde (1% vol/vol final concentration) at room temperature. Cross-linking was stopped with 125 mM glycine for 10 min. Cross-linked chromatin was sonicated to obtain fragments between 250 and 750 base pairs in a Bioruptor Pico sonication device (Diagenode). The sheared chromatin was immunoprecipitated with 1 ug of antibody raised against TERT (Rockland, 600-401-252S) and 0.3 ug STAT5 (Cell Signaling, 94205S). Normal rabbit polyclonal IgG (Cell Signaling, 2729S) was used as a negative control. Negative control was incubated with rabbit IgG and input DNA primary antibody. Chromatin complexes were recovered with ChIP-grade Protein G magnetic beads (9006S, Cell Signaling). DNA was recovered with standard phenol-chloroform extraction. Immunoprecipitated DNA was amplified by quantitative RT-PCR using SYBR green. ChIP primers are listed in Supplementary Table 3.

### Proximity Ligation Assays (PLA)

PLA was performed directly after cell fixation and according to manufacturer instructions. Briefly, cells were blocked with Duolink blocking solution for 60 minutes at 37°C. Samples were then incubated with rabbit TERT and mouse STAT5 antibodies diluted in Duolink Blocking Solution and incubated overnight at 4°C. Samples were then incubated with secondary antibodies conjugated with PLA probes (DUO92002, DUO92004, Sigma) for 2 h at 37°C in a humidity chamber, followed by probe ligation for 30 minutes at 37°C as recommended by the manufacturer. Amplification was performed with Duolink detection kit Red 595 nm for 100 minutes at 37°C (DUO92101, Sigma). Samples were mounted in DAPI-containing mounting media and prepared for image acquisition. PLA foci were quantified using ImageJ relative to the number of cells per field. Fields with over 50 cells and 5 fields per cell line were acquired. Nuclear PLA foci were all puncta circumscribed to the nuclear (DAPI) signal.

### Promoter Analysis

The upstream region of the *RUNX2* (NM_001015051.3) promoter was scanned with LASAGNA and TRANSFAC/ suites.^12,13^ Searches were conducted with data version 2021.3, high-quality matrices, and cut-offs to minimize false positive hits. STAT5 logo analysis was performed with WebLogo (weblogo.berkeley.edu).

### SplitFAST plasmid construction and SplitFAST assay

Plasmids containing full-length human CDS TERT pEZ-M02-CMV-hTERT and STAT5A pEZ-M02_CMV_hSTAT5A were purchased from Genecopeia (catalog numbers EX-Q0450-M02-10 and EX-F0979-M02-10, respectively). Plasmids containing NFAST (pCMV-FRB-NFAST) and CFAST (pCMV-FKBP-CFAST) tags were purchased from The Twinkle Factory (catalog numbers PL058 and PL040, respectively). Fusion plasmids were constructed by PCR assembly. TERT and STAT5A were amplified using primers hTERT-RP-SplitFAST and hTERT-FP-SplitFAST and hSTAT5A-RP-SplitFAST and hSTAT5A-FP-SplitFAST, respectively (Supplementary Table 3). The plasmid pCMV-TERT-NFAST was generated by inserting TERT gene into pAG148-FRB-NFAS using restriction enzymes *Bgl*II and *Kpn*2I. The plasmid pCMV-STAT5A-CFAST was generated by inserting the STAT5A gene into pAG152-FRB-FRB-CFAST11 using restriction enzymes *Bgl*II and *Kpn*2I. Insertions were validated by full plasmid sequencing. pCMV-TERT-NFAST and pCMV-STAT5A-CFAST fusion plasmids were transfected in a 1:1 ratio in control hVICs with XtremeGene 9 DNA transfection reagent (catalog number 39320900, Roche) following manufacturer recommendations. To image the interaction between TERT and STAT5A, cells were imaged in alpha-MEM supplemented with 5 μM μM 4-hydroxy-3,5-dimethoxy benzylidene rhodamine (HBR-3,5DOM, dissolved in DMSO) at d3, d5, and d7 of OST treatment in an EVOS FL microscope (AMA3300, Life Technologies) at 20x with an EVOS Plan Fluor 20x/0.5 objective (AMEP 4698, Life Technologies) at 360/447 nm excitation/emission (DAPI), 470/525 nm excitation/emission (GFP).

### In-Silico Modeling

PDB files for TERT (UniProt 014746) and STAT5A (UniProt K7EIF9) were downloaded from the EMBL-EBI-maintained AlphaFold protein structure database.^14^ The STAT5A file was then uploaded to the HDOCK server as both the receptor and ligand.^15^ The top predicted model for the STAT5A dimer was downloaded and reuploaded as the receptor file, while the TERT file was uploaded as the ligand. The top predicted model was downloaded and visualized with the Mol viewer.^16^ Separately, the residue data were examined to assess which regions of TERT and STAT5 were predicted to localize.

### Animal Model of Aortic Artery Calcification (in vivo calcification)

Vascular calcification was induced in 8-weeks old *Tert^+/+^* and *Tert^-/-^* mice with a 12-week diet regime consisting of chow (LabDiet 5P00, Prolab RMH 3000), supplemented with 0.2% adenine (adenine 0.2%, FD&C Blue No. 2 0.3%, LabDiet 5ZAF) for 6 weeks followed by chow supplemented with 0.2% adenine and 1.8% phosphate (adenine 0.2%, FD&C Blue No. 40 0.3%, sodium phosphate 0.65%, potassium phosphate 1,0%, dicalcium phosphate 3.55%, LabDiet 5ZRX) or 6 weeks.^17^ Both groups included an equal number of male and female mice. At the end of the 12-week diet, aortas were dissected for histological analysis, Von Kossa, and immunohistochemistry.

### Cultured Aortic Segments Calcification (ex vivo calcification)

Aorta were dissected from *Tert^-/-^* and *Tert^+/+^* mice after euthanized with CO2 and cervical dislocation, followed by in situ perfusion with 10 ml of PBS. Collected arteries were cut into 3-4 mm segments and cultured in osteogenic or control media for 14 days in a humidified incubator with 5% CO2 at 37 °C. The medium was changed every 2-3 days. Aortic arteries osteogenic media was prepared as follows: DMEM (11965092, Thermo Fisher Scientific, MA, USA) supplemented with 1% fetal bovine serum (FBS), 1% Antibiotic-Antimycotic (15240062, Thermo Fisher Scientific), 10 mM β-glycerophosphate disodium (G9422, Sigma), 0.25 mM L-ascorbic acid (49752, Sigma), and 10 nM dexamethasone (D4902, Sigma). The control medium was prepared with DMEM supplemented with 1% FBS and 1% Antibiotic-Antimycotic. At the end of the treatments, aortic segments were washed with PBS, fixed in 4% PFA for 4 hours, embedded in paraffin wax, and cut in 10 um slices for histology and immunofluorescence staining.

### Blinding Procedures

Professionally trained technicians handled animal welfare, including weaning and mouse line expansion, diets, sacrifice, and surgery. For experiments involving human samples and in vivo and ex vivo murine experiments, blinding procedures were used. Individuals conducting and analyzing experiments did not know the clinical diagnosis nor treatment groups until after the analysis was performed.

### Sample Size and Power Analysis

The two-sample Cohen’s D algorithm for large-size effects was used.^18^ Power was defined at > 80%, with confidence levels of 95%, alpha at 0.05, and P significance value. The animal size was based on a power analysis that assumes confidence levels of 95%, the probability of finding a statistical difference between groups of 80%, an expected difference of 37% between groups, and a standard deviation of 25%; experiments required at least five mice per group. A 25% redundancy is built-in to allow for mortality associated with treatments and other complications.

### Statistics

Statistical analyses were performed with GraphPad Prism 9.2 (GraphPad Software, San Diego, CA) software. All experiments used at least n = 3 biological replicates and were run in technical duplicates. Statistical comparisons between the two groups were performed by a nonparametric Mann-Whitney *U* test. Statistical comparisons among more than two groups were performed by nonparametric Kruskal-Wallis *H* tests with post hoc Dunn’s multiple pairwise comparisons between groups. Shapiro-Wilk normality test was used to test data distribution. Data are presented as mean ± SD.

---

**Supplemental Figure 1.**
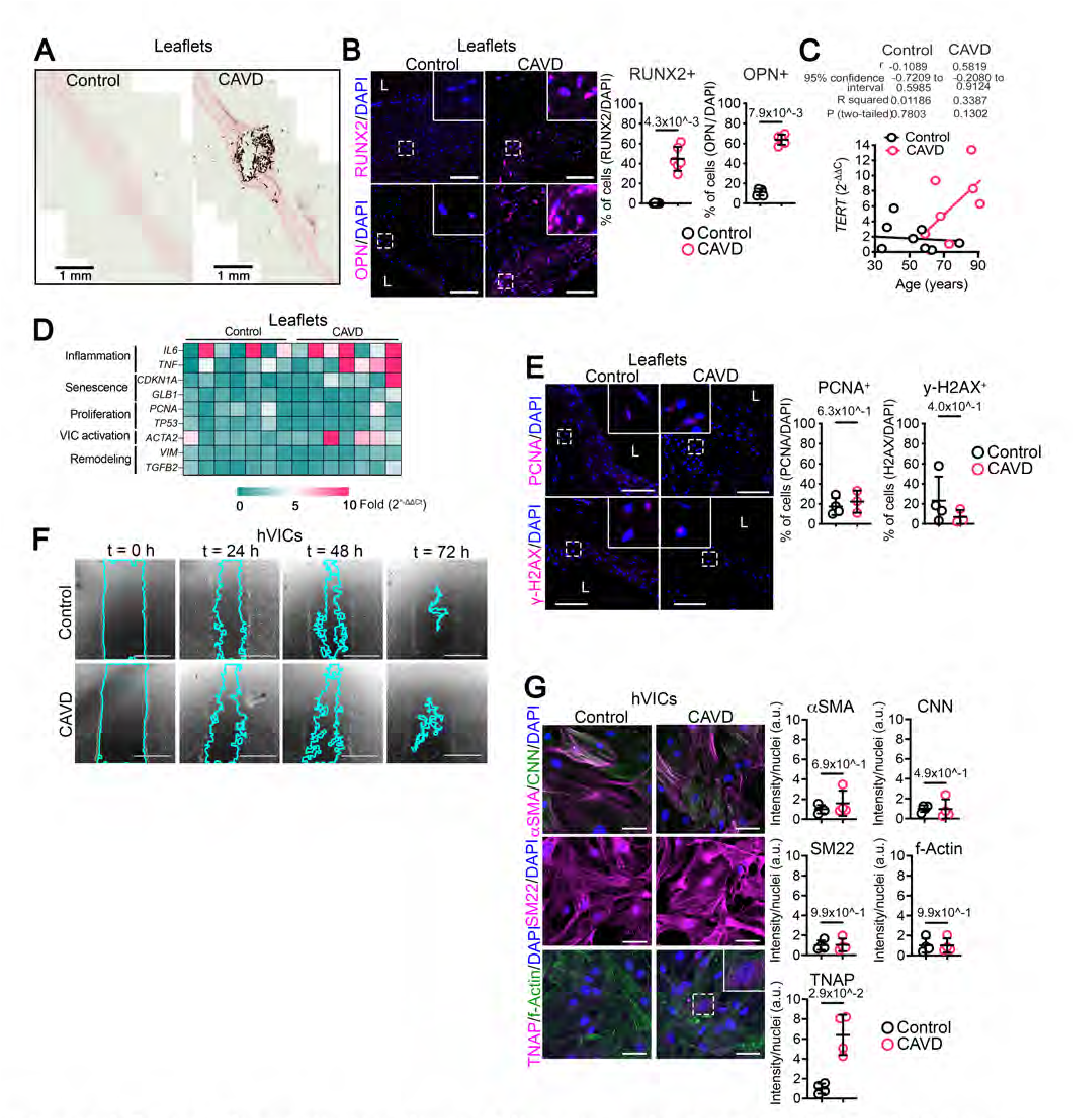
Characterization of human CAVD valves and patient-isolated VICs at baseline. (A) Representative section of Von Kossa staining in control and valve CAVD leaflets tissue. Scalebar 1 mm. (B) Representative serial sections of RUNX2 immunofluorescent staining and OPN in control and CAVD valve tissues. Scalebar 100 μm. Quantification of RUNX2-positive cells and OPN-positive cells on the leaflet specimen is shown on the right graphs. n = 5 control, n = 6 CAVD. (C) Correlation between age of the donor and TERT expression in valve tissues. n = 8 control, n = 9 CAVD. (D) Gene expression profile in the valve leaflets. n = 7 each group. (E) Representative images of immunofluorescent staining of PCNA (top panels) and γ-H2AX (bottom panels) in control and CAVD valve tissues. Scalebar 100 μm. Quantification of PCNA-positive cells and γ-H2AX-positive cells are shown on the right graphs. n = 4 control, n = 3 CAVD. (F) Representative images of a scratch assay in control and CAVD hVICs. Time points are indicated on top of each panel. Blue lines denote the cell migration front. Scalebar 1 mm. n = 10 of each group. (G) Representative images of αSMA, CNN, SM22, TNAP, and f-Actin in control and CAVD hVICs. Scalebar 50 μm. Signal quantification is shown on the right. n = 4 each group. Data are shown as means ± SD. P values were calculated using the Mann-Whitney U test (Figures B, D, E, and G) and the Pearson correlation test (Figure C). L = lumen.

**Supplemental Figure 2.**
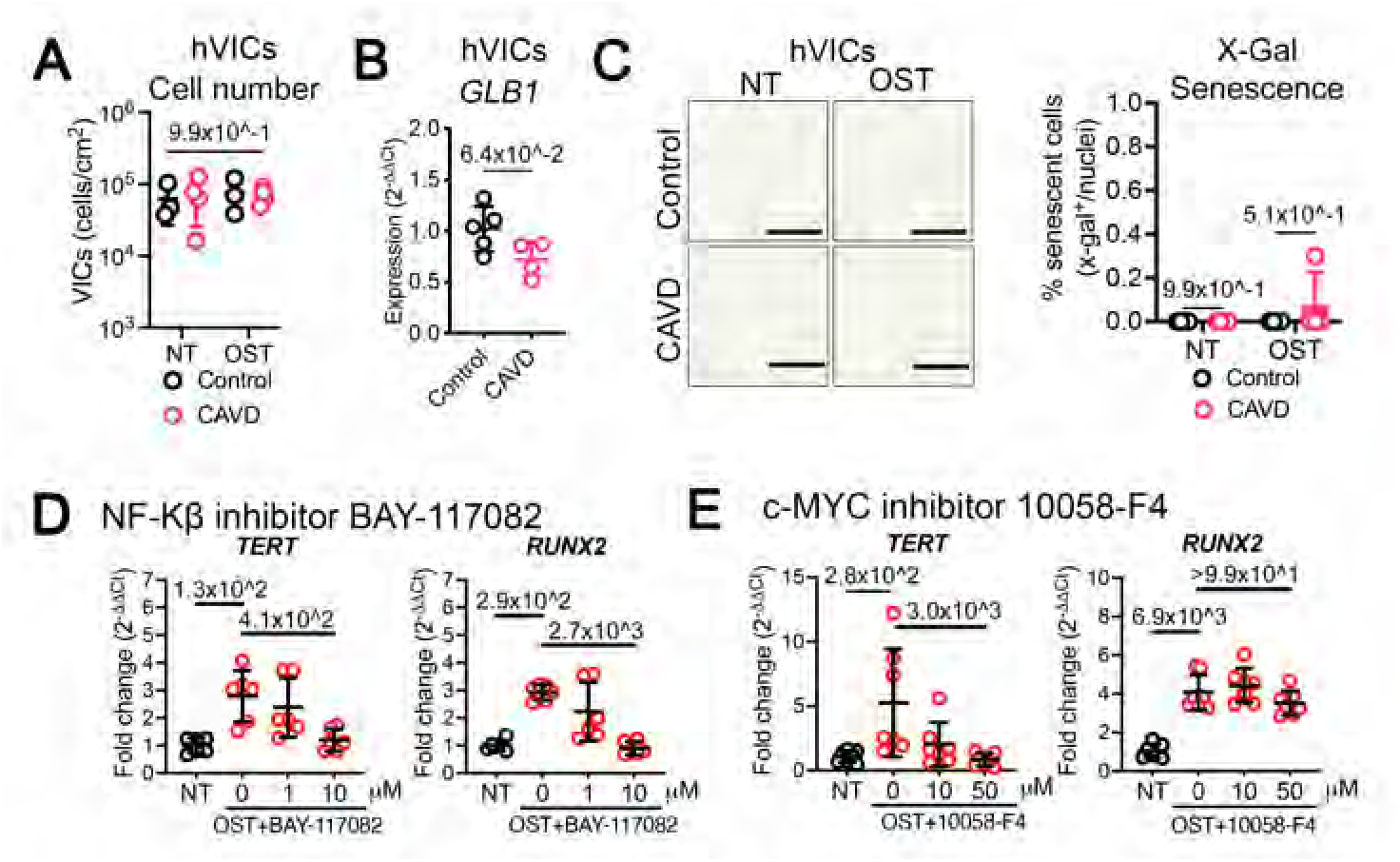
Inhibition of reverse transcriptase activity of TERT does not prevent osteogenic reprogramming of hVICs. (A) Cell counts of control and CAVD hVICs after 14 days of NT or OST stimulation. n = 4 each group. (B) β-galactosidase (BLG1) mRNA transcript quantification in control and CAVD hVICs at baseline. n = 5 control, n= 4 CAVD. (C) Representative images of senescence-associated β-galactosidase activity staining of hVICs after 28 days of NT or OST stimulation. Blue cells were considered positive for senescence. Quantification of senescence is shown on the right graph. n = 4 each group. Scalebar 400 μm. (D) hVICs growing in no treatment (NT), OST media, or OST media supplemented with the NF-κβ inhibitor BAY-117082 for 7 days, n = 6. (E) hVICs growing in no treatment (NT), OST media, or OST media supplemented with the c-MYC inhibitor 10058-F4 for 7 days, n = 7. qRT-PCR quantified TERT and RUNX2 gene expression. In D and E, inhibitors were dissolved in DMSO thus NT and OST were supplemented with the highest volume of vehicle (DMSO) used in treatment groups. Data are shown as means ± SD. P values were calculated with the Mann-Whitney U test (Figure B) and the Kruskal-Wallis H test with Dunn pairwise comparison post hoc test (Figures A, C-E) and shown on top of each graph.

**Supplemental Figure 3.**
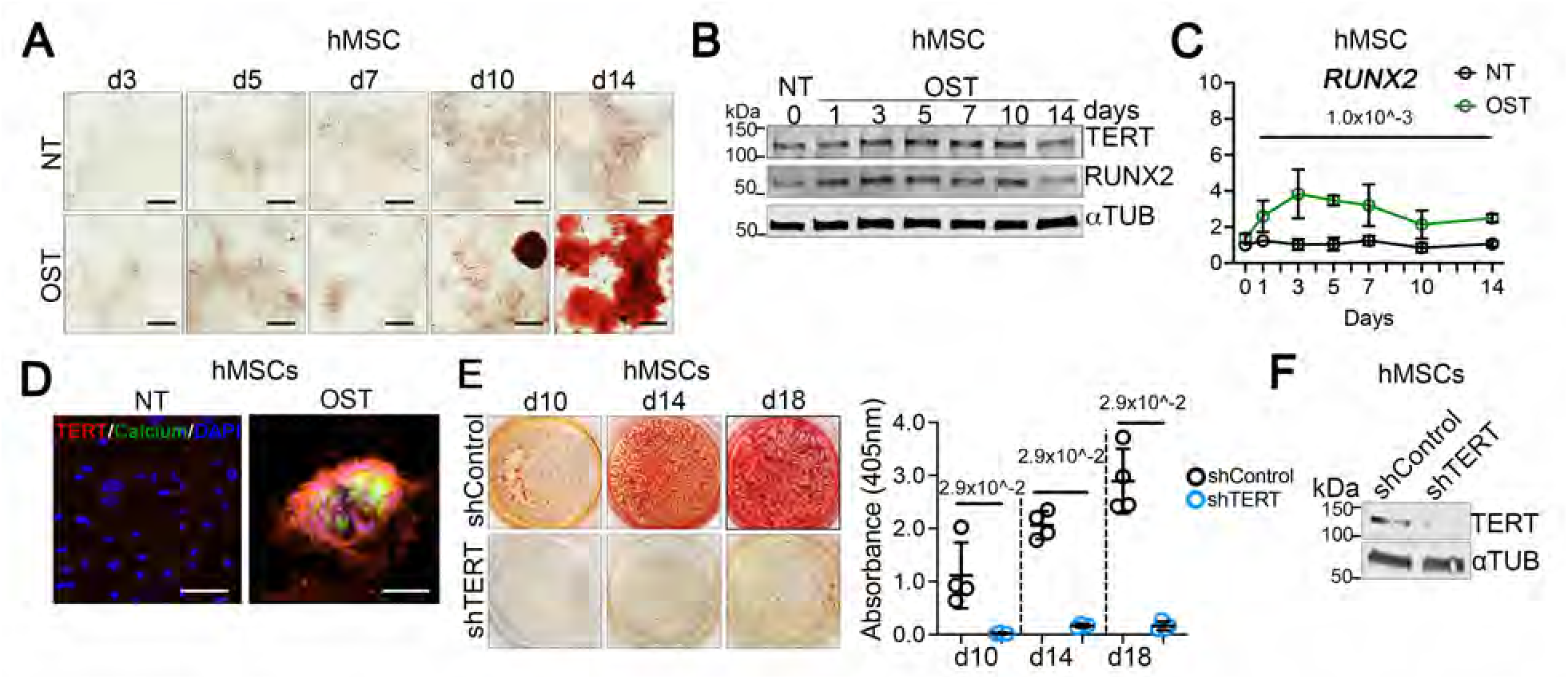
TERT is upregulated and required for the osteogenic differentiation of human mesenchymal stem cells. (A) Representative images of hMSCs stimulated with OST for 14 days. Calcium deposition was visualized by Alizarin Red staining. Scalebar 50 μm. (B) Western blot staining of samples from hMSCs collected during the 14 days of OST treatment. (C) RUNX2 expression profile of differentiating hMSCs stimulated for 14 days with OST media. n = 3. (D) Representative TERT immunofluorescent staining images of hMSCs stimulated with OST for 14 days. Scalebar 100 μm. (E) Representative images of hMSCs infected with shTERT or shControl virus and stimulated with OST for 14 days. Calcium deposition was visualized by Alizarin Red staining. The quantification of calcification is shown on the right graph. n = 3. (F) Western blot staining of samples from hMSCs collected during the 14 days of OST treatment. Data are shown as means ± SD. P values were calculated with the Kruskal-Wallis H test with Dunn pairwise comparison post hoc test (Figure C) and the Mann-Whitney U test (Figure E) and shown on each graph.

**Supplemental Figure 4.**
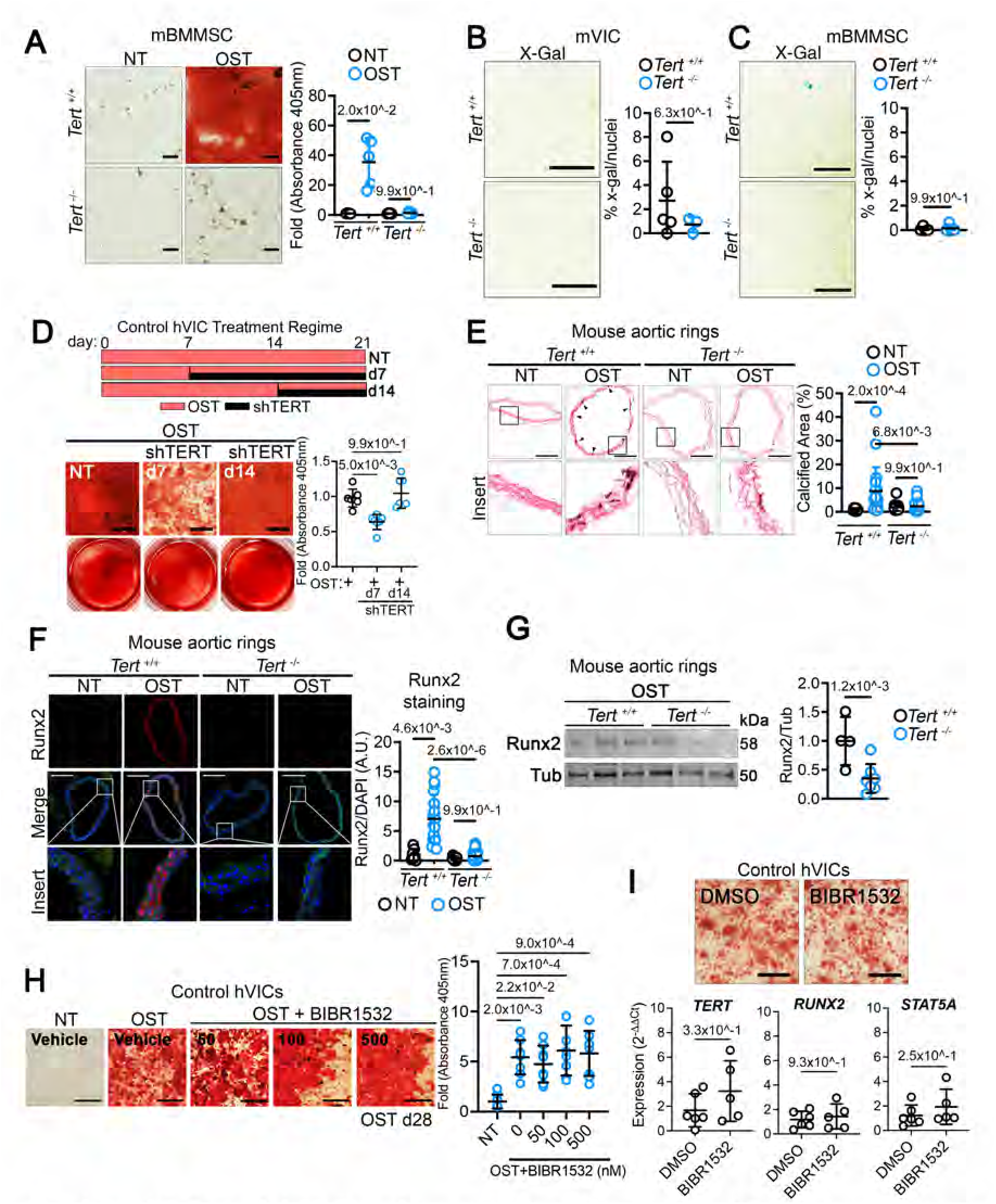
Inhibition of reverse transcriptase activity of TERT does not prevent osteogenic reprogramming of VICs. (A) Representative images of mBMMSCs isolated from *Tert^+/+^* or *Tert^-/-^* mice and stimulated with OST for 21 days. Scalebar 100 μm. Quantification of calcification is presented as average absorbance on the right graphs, n = 5 for each group. (B, C) Representative images of a senescence-associated β-galactosidase (SA-β-gal) staining of mVICs and mBMMSCs isolated from *Tert^+/+^* or F1 *Tert^-/-^* mice after 21 days of OST stimulation. In each group, blue cells were considered positive for senescence, n = 5. (D) Schematic of OST treatment with delayed shTERT transduction (top panel). VICs were transduced with shTERT, and calcification was assessed with Alizarin Red S stain at day 21. Scalebar 400 μm. n = 6 for each group. (E) Representative sections of Von Kossa staining in thoracic aortic rings isolated from *Tert^+/+^* and *Tert^-/-^* mice after 14 days of OST treatment relative to NT control treatment. Quantification is shown on the right. Scale bar 400 µm (F) Immunofluorescence staining of thoracic aortic rings after OST treatment as in B. Quantification of Runx2 is shown on the right graph. Scale bar 400 µm (G) Western blot staining of aortic ring specimens collected and treated in B. Quantification of Runx2 is shown on the right graph. (H) Human VICs were treated with the TERT inhibitor BiBR1532 and stimulated with OST, and calcification was assessed on day 28, with n = 5 in each group. (I) TERT, RUNX2, and STAT5A expression were evaluated on day 14, n = 6 DMSO, n = 5 BIBR1532. Data are shown as means ± SD. P values were calculated with the Kruskal-Wallis H test with Dunn pairwise comparison post hoc test (Figures A, B, C, E, H) and the Mann-Whitney U test (Figures D, F, G, I) and are shown on top of each graph.

**Supplemental Figure 5.**
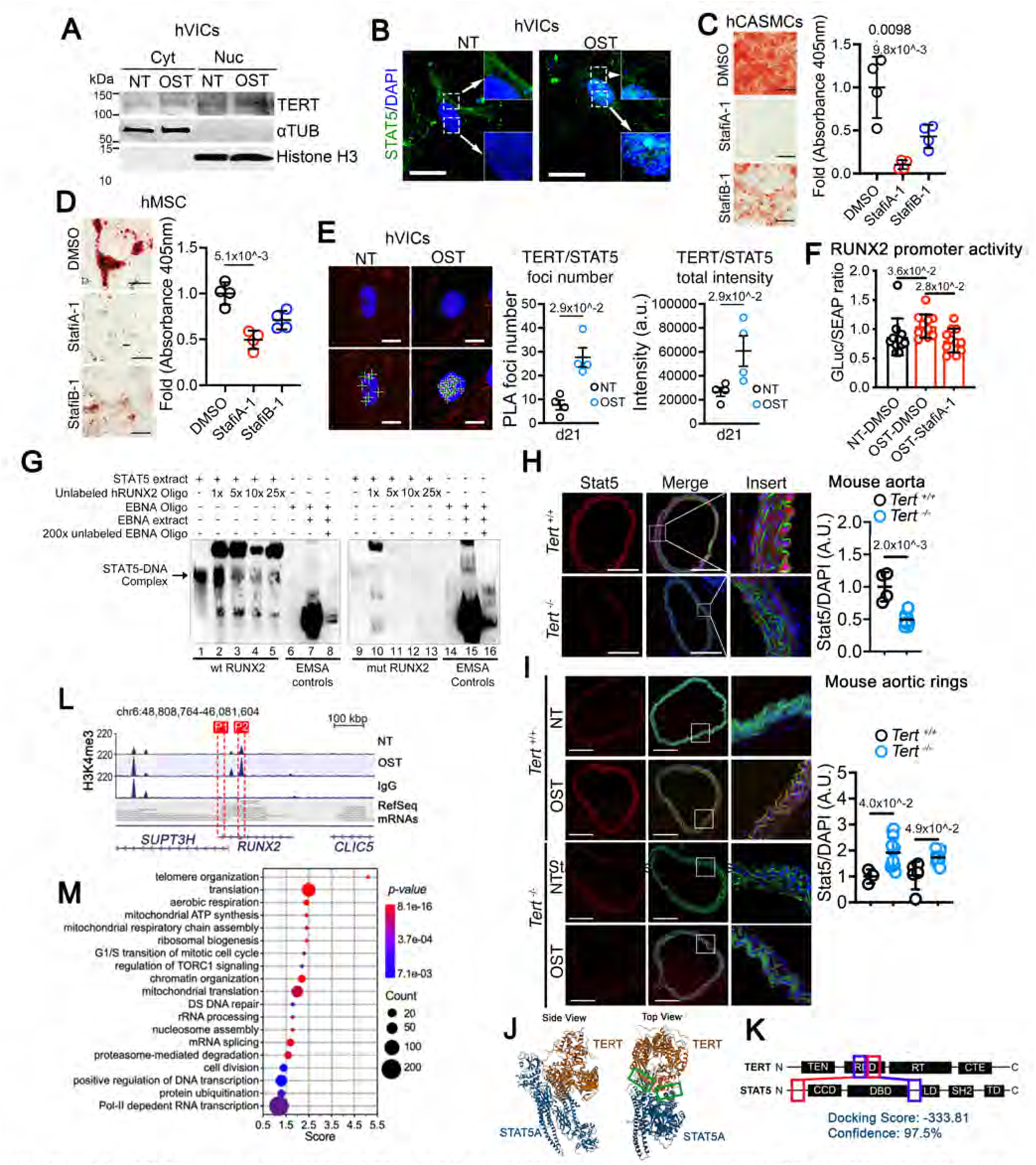
STAT5 is upregulated, binds to TERT, and is required for osteogenic reprogramming of VICs. (A) Western blot analysis of cytoplasmic and nuclear fractions of TERT from control VICs obtained after seven days of OST stimulation. TUB, (tubuline); H3, (Histone H3); Cyt, cytoplasmic fraction; Nuc, nuclear fraction. (B) Representative immunofluorescence image of STAT5 subcellular distribution after 14 days of NT and OST stimulation. Scalebar 40 μm. (C) Representative images of human CASMCs treated with 10 μM of the STAT5 inhibitors StafiA-1 or StafiB-1 during the 14 days of OST stimulation. Scalebar 400 μm. n = 4 for each group. (D) Representative images of human MSCs treated with 10 μM of the STAT5 inhibitors StafiA-1 or StafiB-1 during the 21 days of OST stimulation. Scalebar 400 μm. n = 4 for each group. (E) Representative images of TERT/STAT5 complex (red foci) detected by proximity ligation assay (PLA) in the nucleus of control VICs. Scalebar 25 μm. n = 4 for each group. (F) Relative luciferase activity in hVICS after transfection of *hRUNX2* Promoter GLuc-ON pEZX-PG04.1. Samples were analyzed 72 hours after OST treatment. STAT5A inhibitor StafiA-1 (10 μM) was dissolved in DMSO. NT and OST conditions were also supplemented with DMSO, n = 10 per group. (G) Biotin-labeled 60 bp oligos containing the −194 STAT5 binding sequence on the promoter of human RUNX2 were incubated cell extracts from HEK293s in which STAT5 protein was overexpressed (lane 1). Extracts were also incubated with an increasing amounts of unlabeled oligos (lanes 2-5). Cell extract was also incubated with a oligo containing a mutated STAT5A site (mut RUNX2 DNA) (line 9) and incubated with an increased amount of unlabeled mut RUNX2 oligo (lanes 10-13). Blots include the positive and negative controls include in the LightShift Chemiluminescent EMSA Kit (lanes 6-8 and 14-16). (H) Representative immunofluorescent images of Stat5 expression after 12 weeks of high adenine high phosphate diet on Tert-/- and Tert-/-mice aorta. Scale bar 400 µm. Quantification of Stat5 expression is shown on the right panel. (I) Representative images of Stat5 immunofluorescence staining of thoracic aortic rings isolated from Tert+/+ and Tert-/-mice after 14 days of OST treatment relative to NT control treatment. Quantification of Stat5 is shown on the right graph. Scale bar 400 µm. (J) Mol*3D Viewer rendering of the TERT-STAT5 dimer interaction predicted by the HDock. Red boxes indicate the position of the interaction between TERT and STAT5. (K) The diagram shows the regions involved in the predicted interaction as indicated in A, and the docking and confidence scores predicted by HDock are indicated. TEN: DNA anchor domain for processive telomeric repeat addition, RBD: Telomerase RNA binding domain, RT: Finger and palm region of the reverse transcriptase domain, CTE: C-terminal extension or putative thumb domain of the reverse transcriptase, CCD: Coiled-coil domain, required for nuclear localization, DBD: DNA binding domain, HLK: Helical linker and DNA binding domain, SH2: Domain required for activation and dimerization, TD: Transactivation domain. (L) H3K4me3 CUT&Run sequencing tracks in genomic regions of Runx2 in no-treatment control and osteogenic condition-treated human VICs. (M) Top 20 GO pathways enriched in genes with significant loss of H3K4me3 in OST-vs NT-treated human VICs. Data are shown as means ± SD. P values were calculated with the Mann-Whitney U test (Figures E and G) and the Kruskal-Wallis H test with Dunn pairwise comparison post-test (Figures C, D, and H) and indicated in each graph.

**Supplemental Figure 6.**
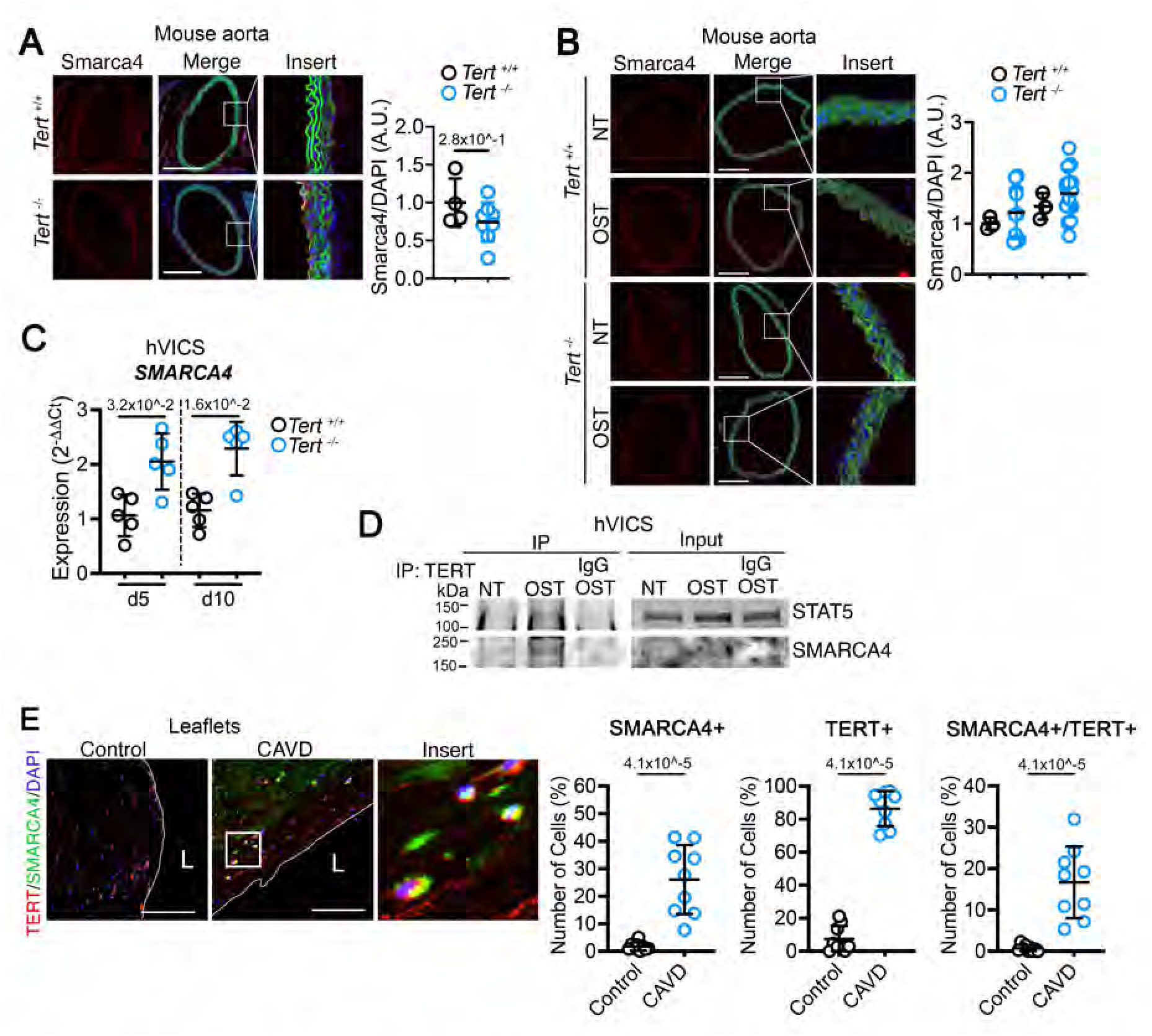
Chromatin remodeling enzyme SMARCA4 is upregulated and interacts with TERT during the osteogenic reprogramming of VICs. (A) Representative immunofluorescent images of Smarca4 expression after 12 weeks of high adenine high phosphate diet on *Tert^-/-^* and *Tert^-/-^* mice aorta. Scale bar 400 µm. Quantification of Smarca4 expression is shown on the right panel. (B) Representative images of Smarca4 immunofluorescence staining of thoracic aortic rings isolated from *Tert^+/+^* and *Tert^-/-^* mice after 14 days of OST treatment relative to NT control treatment. Quantification of Smarca4 is shown on the right graph. (C) SMARCA4 expression was evaluated on days 5 and 10 on VICs stimulated with OST. n = 5 each group. (E) TERT/STAT and TERT/SMARCA4 interaction was detected by co-immunoprecipitation upon osteogenic stimulation on VIC. (E) Representative TERT and SMARCA4 immunofluorescent staining in control and CAVD tissues. The number of positive cells is shown on the right graphs. n = 9 control, n = 9 CAVD. Scalebar 100 μm. Data are shown as means ± SD, and P values were calculated with the Mann-Whitney U test and indicated on each graph in Figures A and C. L = lumen.

**Supplemental Table 1:** Patient information.

| Patient ID | Age | Sex | Valve | Cause of Death | Calcification<br>(Von Kossa Staining) | Hypertension | Diabetes | Smoking | Alcohol |
| --- | --- | --- | --- | --- | --- | --- | --- | --- | --- |
| 2016-034 | 91 | Male | Tricuspid | Natural causes | Yes | No | No | No | Unknown |
| 2016-037 | 31 | Male | Tricuspid | Brain anoxia due to cardiovascular complications | No | No | No | Unknown | Yes |
| 2016-038 | 81 | Male | Tricuspid | Myocardial Infarction | No | No | No | Unknown | Unknown |
| 2016-042 | 22 | Female | Tricuspid | Brain anoxia due to head trauma | No | No | No | No | Unknown |
| 2016-045 | 59 | Female | Tricuspid | Stroke | No | PH | Yes | Yes | Unknown |
| 2016-046 | 65 | Male | Tricuspid | Head trauma from accident | No | No | No | Unknown | Unknown |
| 2016-048 | 56 | Female | Tricuspid | Stroke | No | HTN | No | Unknown | Unknown |
| 2016-050 | 65 | Male | Tricuspid | Stroke | No | HTN | No | Unknown | Yes |
| 2016-051 | 65 | Male | Tricuspid | Natural causes | Yes | HTN | No | Yes | Yes |
| 2016-053 | 73 | Male | Tricuspid | Congestive heart failure | Yes | No | No | No | Unknown |
| 2016-054 | 63 | Female | Tricuspid | Hemorrhage and Rupture AAA | No | No | Yes | No | Unknown |
| 2016-060 | 73 | Male | Tricuspid | Cardiac arrest | Yes | No | No | Unknown | Unknown |
| 2016-061 | 67 | Male | Tricuspid | Natural causes | Yes | No | Yes | Unknown | Unknown |
| 2016-062 | 59 | Male | Tricuspid | Cardiac arrest | Yes | No | No | Yes | Unknown |
| 2016-065 | 58 | Male | Tricuspid | Stroke | No | HTN | Yes | Yes | Unknown |
| 2016-068 | 68 | Female | Tricuspid | Intracranial hemorrhage | Yes | No | No | Yes | Unknown |
| 2016-072 | 62 | Female | Tricuspid | Cardiac arrest | Yes | No | No | No | Unknown |
| 2016-073 | 41 | Female | Tricuspid | Traumatic arrest and motor vehicle accident | No | HTN | No | Yes | Unknown |
| 2016-078 | 34 | Female | Tricuspid | Heart Failure | No | No | No | Unknown | Unknown |
| 2016-079 | 59 | Male | Tricuspid | Respiratory failure | No | HTN | No | No | Unknown |
| 2016-083 | 52 | Male | Tricuspid | Stroke | No | PAH | No | Yes | Unknown |
| 2016-085 | 52 | Male | Tricuspid | Myocardial Infarction | No | PH | No | Unknown | Unknown |
| 2016-091 | 59 | Female | Tricuspid | Stroke | No | HTN | No | Unknown | Unknown |
| 2016-092 | 47 | Female | Tricuspid | Stroke | No | No | No | No | Unknown |
| 2016-093 | 43 | Male | Tricuspid | Cardiac arrest | Yes | HTN | Yes | No | Unknown |
| 2016-095 | 87 | Male | Tricuspid | Myocardial Infarction | Yes | HTN | No | Yes | Unknown |
| 2016-098 | 57 | Male | Tricuspid | Cardiac arrest | No | HTN | No | Yes | Unknown |
| 2016-101 | 37 | Female | Tricuspid | Cardiac Death | No | No | No | Unknown | Unknown |
| 2016-102 | 58 | Female | Tricuspid | Cardiac arrest and asystole | Yes | No | No | Yes | Unknown |
| 2016-103 | 72 | Male | Tricuspid | Cardiac arrest | Yes | HTN | No | Unknown | Unknown |
| 2016-105 | 86 | Male | Tricuspid | Cardiac arrest | Yes | No | No | No | Unknown |
| 2016-118 | 62 | Male | Tricuspid | Cardiac arrest | Yes | No | No | No | Unknown |
| 2016-146 | 79 | Male | Tricuspid | Cardiac arrest | No | No | No | Yes | Unknown |
| 2017-005 | 66 | Female | Tricuspid | Stroke | No | No | No | Unknown | Unknown |
| 2017-025 | 57 | Female | Tricuspid | Brain anoxia due to choking | Yes | No | Yes<br>Type II | Unknown | Unknown |
| 2017-035 | 48 | Female | Tricuspid | Brain anoxia due to CO poisoning | No | No | No | Unknown | Unknown |
| 2017-051 | 56 | Male | Tricuspid | Brain anoxia due to cardiac arrest | No | No | No | Unknown | Unknown |
| 2017-069 | 37 | Male | Tricuspid | Motor vehicle accident | No | No | No | Unknown | Unknown |
| 2017-076 | 68 | Female | Tricuspid | Myocardial Infarction | Yes | No | No | No | Unknown |
| 03-0127 | 61 | Male | Tricuspid | Surgical replacement | No | HTN | No | Unknown | Unknown |
| 03-0129 | 68 | Male | Tricuspid | Surgical replacement | No | HTN | Yes<br>Type II | Unknown | Unknown |

HTN = Hypertension
PH = Pulmonary Hypertension
PAH = Pulmonary Arterial Hypertension

**Supplemental Table 2:**
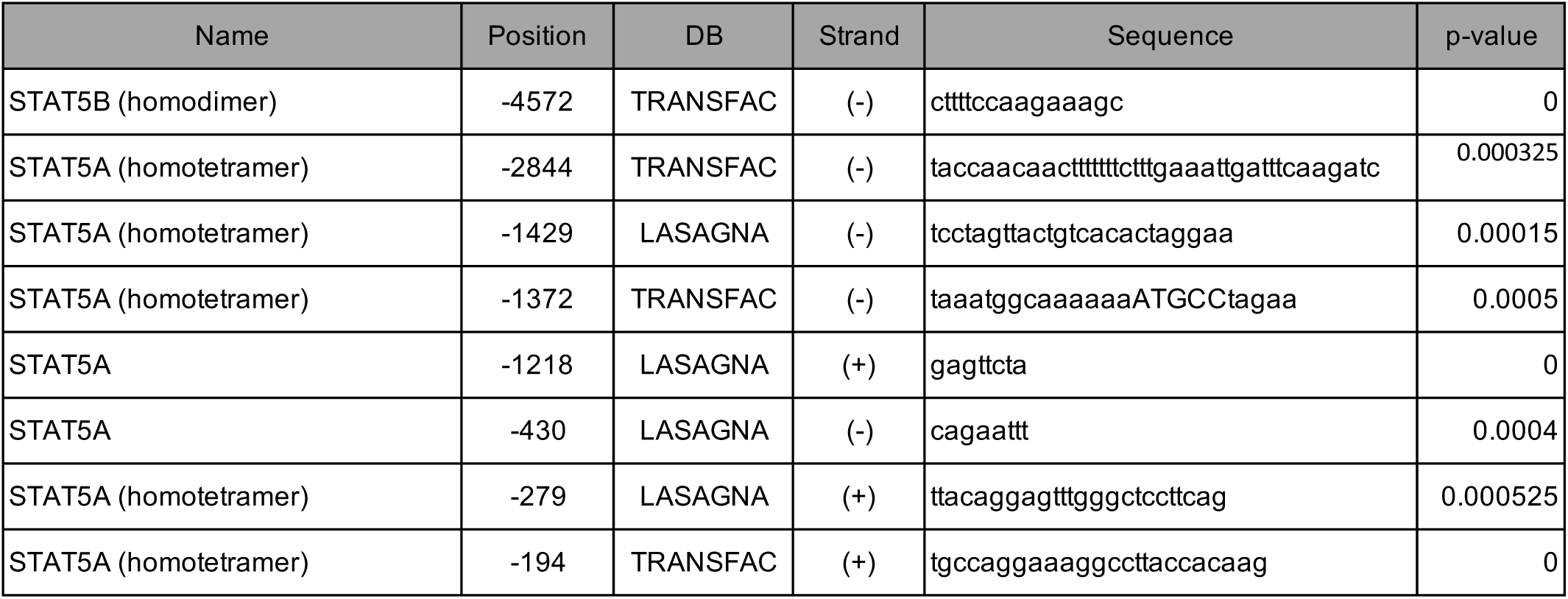
LASAGNA and TRANSFAC STAT5 Search on RUNX promoter.

**Supplemental Table 3:**
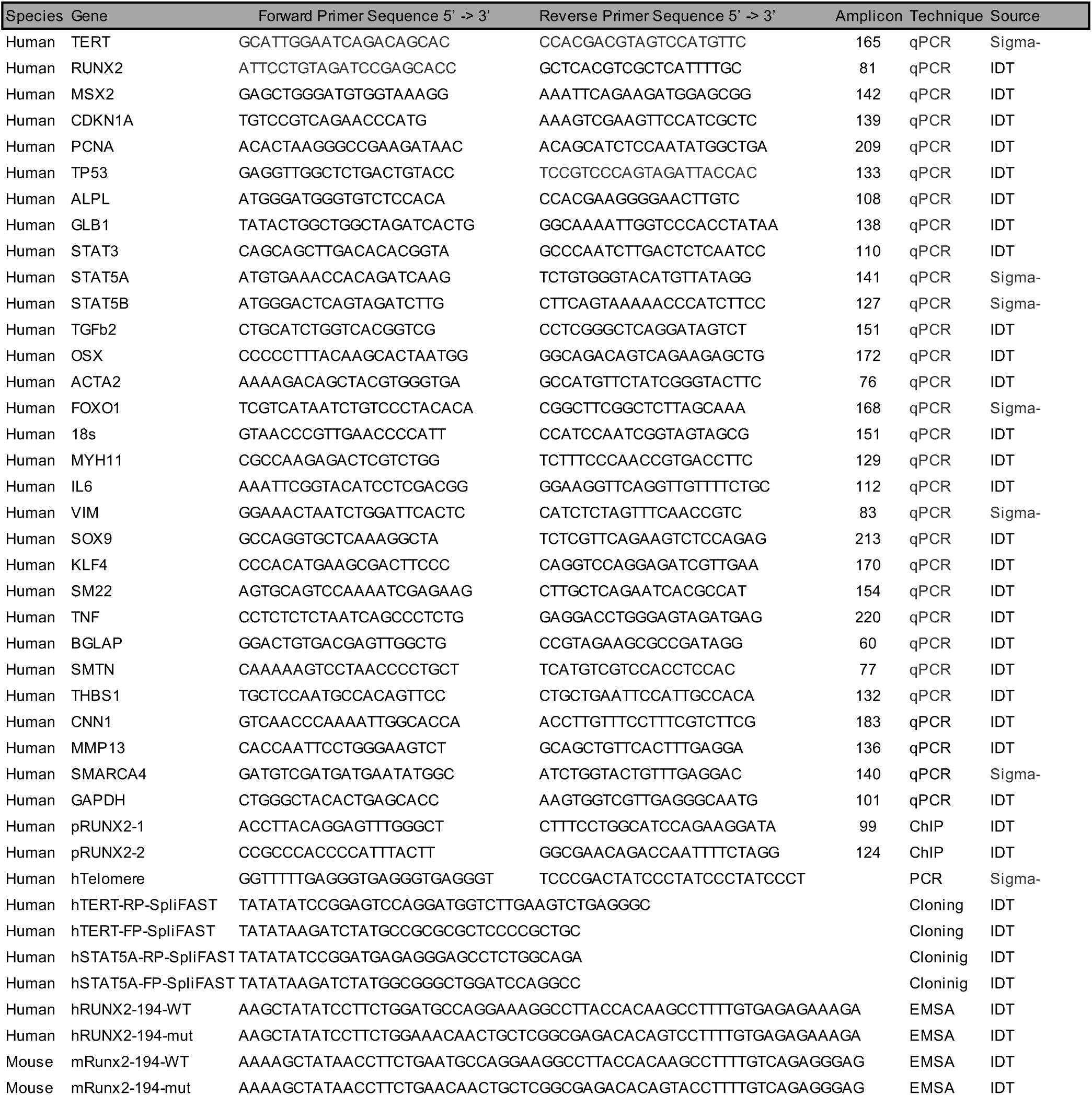
qPCR, Cloning Primers, and Oligos.

